# *Pseudomonas aeruginosa* balances cytotoxicity and motility to counter phagocytosis by macrophages

**DOI:** 10.64898/2026.04.02.716043

**Authors:** Tania Distler, Chia-Ni Tsai, Aaron Weimann, Zainebe Al-Mayyah, Lucas A. Meirelles, R. Andres Floto, Alexandre Persat

## Abstract

During chronic lung infections, *Pseudomonas aeruginosa* diversifies under selection from antibiotics, metabolic constraints, and host defenses. Macrophages are key sentinels of the innate immune system and play a central role in clearing airway pathogens. Yet, how they process heterogeneous bacterial populations remains poorly understood. Here, we investigate how *P. aeruginosa* evades phagocytosis under conditions that mimic chronic infection. We use an attenuated mutant lacking a functional type III secretion system (T3SS), which reduces macrophage killing, allowing us to isolate determinants of bacterial susceptibility to phagocytosis. Using transposon insertion sequencing (Tn-seq), we identify bacterial fitness factors under phagocytic selection. Our screen reveals that disruption of genes involved in swimming and twitching motility reduces uptake by macrophages. We find that motility defects interfere with the physical interactions between bacteria and macrophages. Live-cell imaging shows that motility-deficient bacteria exhibit reduced surface exploration and unstable attachment to macrophages, limiting their internalization. Clinical isolates with reduced swimming or twitching motility display similarly impaired uptake. Restoring T3SS activity in these motility mutants rescues cytotoxicity toward macrophages, with one notable exception: flagellum-less, hyper-piliated *P. aeruginosa* remains avirulent and resistant to phagocytosis due to their lack of engagement with macrophages. Together, these results support two distinct immune evasion strategies: during chronic infection, reduced motility promotes a “freeze”-like state that limits detection and engulfment, whereas during acute infection, *P. aeruginosa* adopts a “fight”-like strategy by activating its T3SS to eliminate macrophages.

**Summary:** How immune cells recognize and eliminate bacteria is typically explained by molecular signaling, yet the role of physical interactions remains unclear. We show that bacterial motility is a key determinant of phagocytosis by macrophages. Using functional genomics and live imaging, we find that swimming and twitching motility promote bacterial uptake by enabling effective surface exploration and stable physical engagement with macrophages. Loss of motility commonly observed in chronic *P. aeruginosa* infections reduces these interactions and allows bacteria to evade engulfment. In contrast, during acute infection, bacteria rely on T3SS-mediated killing of macrophages, independently of motility. These findings reveal that phagocytosis is governed not only by immune recognition but also by bacterial mechanical behavior, and identify a shift in host-pathogen interactions associated with chronic versus acute infection. More broadly, this work establishes mechanics as an important dimension of immune evasion.

## Introduction

*P. aeruginosa* is a leading cause of healthcare-associated infections and listed as a high priority pathogen for the development of new antimicrobial therapies[1]. This opportunistic pathogen readily infects individuals with chronic respiratory disease, including chronic obstructive pulmonary disease (COPD) and cystic fibrosis (CF), as well as mechanically ventilated patients in intensive care[2]. In chronic lung infections, *P. aeruginosa* colonizes its host for years, experiencing immune attacks and antibiotics which shape the evolution of genotypically and phenotypically diverse populations. A single chronically infected patient can thus be colonized by *P. aeruginosa* clones of various motilities, metabolism and abilities to form biofilms[3,4]. While the vast majority of *P. aeruginosa* acute infection isolates are highly cytotoxic and cause drastic tissue damage, isolates from chronic infection typically show significantly decreased virulence[5]. Most of these isolates have defects in their ability to secrete effectors of the type III secretion system (T3SS), the main cytotoxicity factor in *P. aeruginosa*[6,7]. Mutations that silence transcriptional activators of T3SS (*exsA*) or deletion of entire gene clusters are typical causes of loss of T3SS function[8,9].

Tissue-resident macrophages represent one of the first lines of defense against microbial invaders. In addition to phagocytosis, macrophages orchestrate inflammatory signaling, recruit other immune effectors to fight microbial invaders, and contribute to tissue homeostasis and repair during infection resolution[10–12]. Macrophages engage with bacteria via surface receptors that either directly bind to microbial surface motifs or bridging antibodies and complement proteins, which initiates actin-driven engulfment and lysosomal degradation[13,14]. In the lung, these multifaceted macrophage functions play a central role in responding to and clearing *P. aeruginosa* infections[15,16]. Yet, despite being sampled in abundance from patient sputum[15,17], it remains unclear how they counteract diversified *P. aeruginosa* phenotypes found in chronic infections. In addition, knowledge about the specific involvement of tissue-resident and recruited macrophage populations in clearing *P. aeruginosa* infections is still very limited[18].

During airway colonization, pathogens must cope with macrophages and neutrophils at the mucosal surface. *P. aeruginosa* employs a rather offensive strategy to repel phagocytotic cell attacks: by engaging its T3SS, it directly kills these predators after physical contact[19]. Given the frequent loss of T3SS function during chronic infection, *P. aeruginosa* must employ alternative survival strategies. Biofilm formation, a typical adaptation in chronic infections providing protection against antibiotics, can serve as a shield against phagocytes[20]. Both increased size of bacterial aggregates and elevated production of extracellular polymeric substance (EPS) at the single-cell level can impede engulfment and further modulate inflammatory responses of immune cells[21–24]. Yet, how *P. aeruginosa* evolution during chronic infection broadly impacts its sensitivity to phagocytosis remains unresolved.

*P. aeruginosa* rotates a single polar flagellum to drive swimming in liquid, while extension-retraction cycles of type IV pili (T4P) power twitching motility on surfaces. *P. aeruginosa* clones sampled from chronic infection are often non-motile[25–27]. Loss of flagellar expression causes phagocytic evasion, indicating effects on bacterial adhesion or downstream signaling induced by flagellin[28]. Indeed, flagellin-mediated TLR5 activation enhances bacterial uptake and killing by alveolar macrophages[29]. However, deletions in stator proteins required for flagellar rotation also reduce phagocytic uptake[30,31]. While T4P have been implicated in non-opsonic engulfment of *P. aeruginosa* by macrophages and neutrophils[32,33], mechanistic understanding is still lacking. In addition, phagocytosis studies are predominantly conducted in highly virulent *P. aeruginosa* strains modeling acute infections. But how motility influences phagocytosis under chronic conditions is largely unexplored.

To address these questions, we investigated *P. aeruginosa* selection against macrophage uptake in a chronic evolution context. We hypothesized that phagocytic evasion strategies are more critical in chronic strains than acute conditions, due to improved immune cell survival. We applied functional genomics to map fitness determinants during macrophage infection with an avirulent, chronic-like *P. aeruginosa* mutant. Guided by these results, we then used live imaging to examine in detail how bacterial motility modulates phagocytic uptake. This approach also allowed us to identify the relative contribution of motility systems in bacteria-macrophage competition between chronic and acute conditions.

### A functional genomics screen identifies bacterial factors influencing phagocytosis

To identify *P. aeruginosa* determinants of survival during phagocytosis, we implemented a transposon sequencing (Tn-seq) screen[34] in macrophages. We used the THP-1 human monocytic cell line due to its reproducibility and scalability compared to primary monocytes. We first assessed the cytotoxicity of the lab strain PAO1 towards THP-1 macrophages by monitoring cell death with the DRAQ7 dye. At MOI 10, *P. aeruginosa* killed most macrophages. The killing phase started around 2 h post-infection and only 23 % of macrophages survived at 10 h post-infection. Rapid macrophage killing confounds factors that influence phagocytosis, thereby complicating the interpretation of Tn-seq experiments. To circumvent these limitations and to better emulate phagocytosis of chronic *P. aeruginosa* isolates, we tested the PAO1 Δ*popB* strain mimicking T3SS loss of function mutations commonly found in chronic infections[6,7]. We confirmed that THP-1 macrophages showed improved survival when infected with Δ*popB* relative to WT (46 % of cells remained alive 10 h post-infection). Δ*popB* uptake by THP-1 macrophages furthermore increased 2-fold compared to WT as indicated by CFU quantifications after gentamycin exclusion assay. We concluded that attenuated virulence constitutes suitable conditions to perform a functional genomics screen. We thus constructed the Tn5 transposon (Tn) library in the Δ*popB* PAO1 background. We confirmed that the Tn library mirrors Δ*popB* cytotoxicity and phagocytosis (Fig. 1a,b, S1).

**Figure 1:**
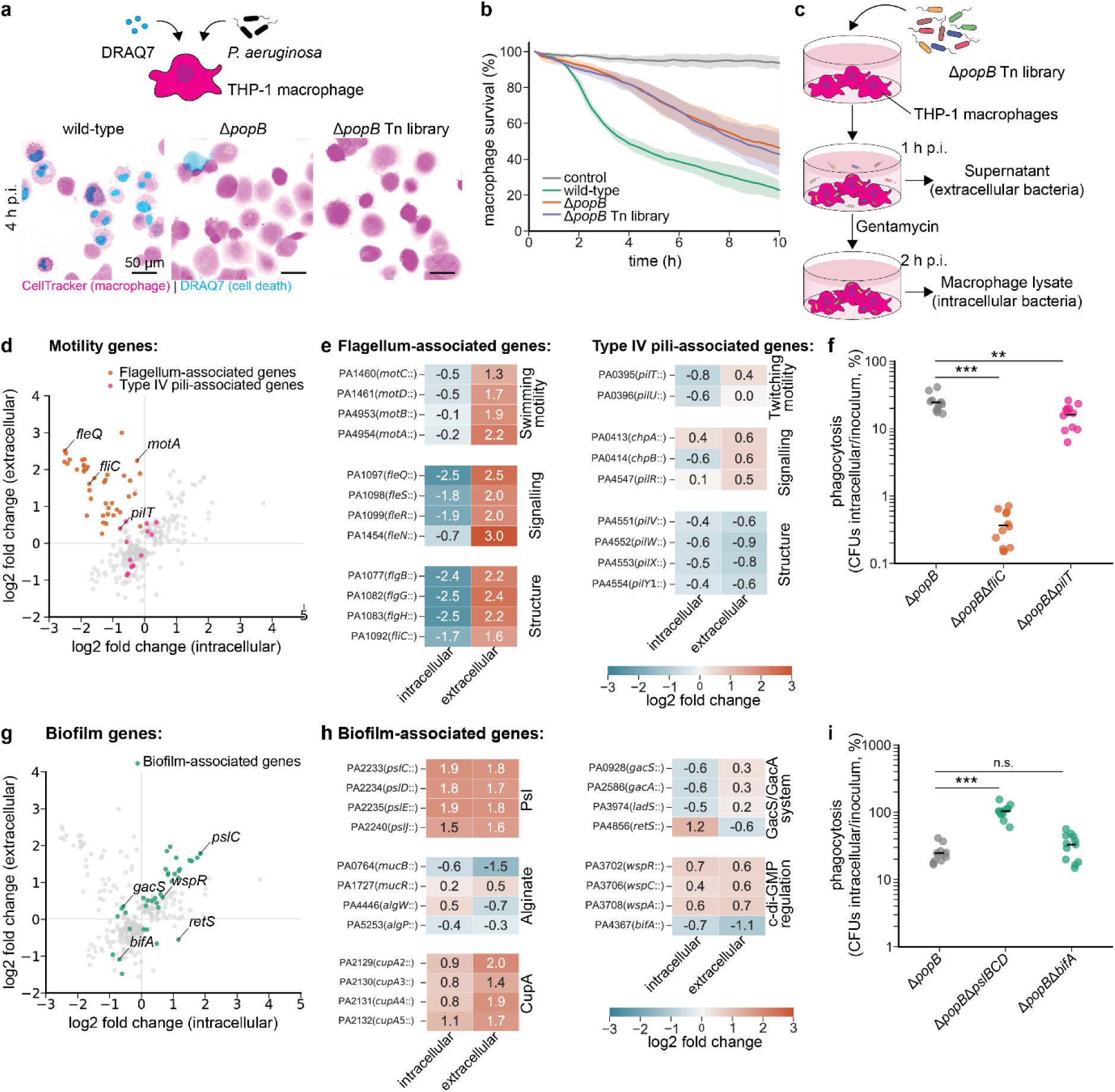
Tn-seq reveals *P. aeruginosa* selection during phagocytosis. **a**, Schematic and exemplary images of the cytotoxicity assay. THP-1 macrophages stained with CellTracker were infected with *P. aeruginosa* and co-incubated with the dye DRAQ7 marking cell death. The cytotoxicity of the PAO1 wild-type strain was compared to the T3SS mutant Δ*popB* and a Transposon (Tn) library constructed in the Δ*popB* background. Images represent maximum z-projections across 5 planes (2 µm step size). **b**, Macrophage survival, defined as the percentage of DRAQ7-negative cells, was quantified over 10 h of infection. Uninfected macrophages served as a control. Thick line, mean; shaded area, standard deviation across biological replicates (n=3). **c**, Depiction of the Tn-seq experimental design. The Tn library was added to THP-1 macrophages at MOI 10. 1 h post-infection (p.i.) the supernatant was harvested followed by the removal of extracellular bacteria by gentamycin for 1 h. Intracellular bacteria were subsequently retrieved by macrophage lysis. Fitness effects were determined in extracellular and intracellular fractions compared to the inoculum. **d**,**g**, Correlation of fold changes between extra- and intracellular fractions highlighting genes that regulate bacterial motility including flagellum- and T4P-associated genes (d) as well as genes that are involved in biofilm formation (g). Only genes with significant fold changes (adjusted p-value<0.05) in at least one fraction are shown. **e**,**h**, Fold changes of transposon insertions in representative genes linked to flagellar and T4P function (e) or biofilm formation (h) are presented for intracellular and extracellular fractions. **f**,**i**, CFU quantification of bacterial uptake by THP-1 macrophages at MOI 10 using clean deletion mutants to validate the Tn-seq results. Among the motility genes, Δ*popB*Δ*fliC* deficient in swimming and Δ*popB*Δ*pilT* lacking twitching motility were tested (f). As representative mutants for biofilm regulation, Δ*popB*Δ*pslBCD*, deficient in Psl biosynthesis, and Δ*popB*Δ*bifA*, exhibiting accelerated biofilm formation due to high c-di-GMP levels, were chosen (i). Black line, mean; circles, technical replicates (from 3 independent experiments). Statistics were assessed by one-way ANOVA with Dunnett’s post-hoc test.

We incubated the Δ*popB* Tn library with THP-1 macrophages for 1 h at MOI 10 and collected the supernatant for sequencing. We next cleared the remaining extracellular bacteria by incubating macrophages with gentamycin. We recovered the surviving intracellular bacterial population by lysing macrophages to obtain a second output sample. We applied an additional outgrowth step to bacterial libraries from the inoculum, supernatant, and macrophage lysates and sequenced these samples to measure the fitness changes of mutations relative to the inoculum (Fig. 1c).

By comparing the intracellular populations to the inoculum, we identified 90 genes whose disruption correlated with intracellular enrichment. These predominantly included mediators of biofilm formation. Mutants in polysaccharide biosynthesis and positive regulators of their secretion were more prevalent intracellularly. Conversely, mutants in 145 genes showed decreased intracellular fitness. These were strongly associated with flagellar motility and assembly, cell wall organization, and type IV pili assembly and function (Table S4, S7, Fig. S2).

To discriminate mutations that affect bacterial uptake during phagocytosis from those that broadly influence survival in the presence of macrophages, we compared data from the supernatant and intracellular populations. Mutants with enhanced uptake were depleted from the supernatant and enriched intracellularly. We identified 77 genes with anticorrelated fitness between fractions indicating that those modulate phagocytosis, the 282 other genes affected overall bacterial survival as their intra- and extracellular fitness was positively correlated (Table S5).

This dataset highlighted a strong prevalence for mutations in genes involved in *P. aeruginosa* motility and biofilm formation. Flagellar mutants, disrupting genes coding for structural components (e.g., *fliC),* stators driving flagellar movement (e.g., *motA*, *motC*) and regulators of flagellar expression (e.g., *fleQ*[35]), were uniformly enriched in the supernatant compared to the intracellular environment. Meanwhile, the effects of mutations in T4P genes were more nuanced: transposon insertions in the retraction motor gene *pilT* or the associated mechanosensory signaling gene *chpB* [36,37] mildly reduced uptake by macrophages, although mutations in other pilus-associated loci did not follow this pattern (Fig. 1d,e). To validate the roles of swimming and twitching motility in modulating phagocytosis, we constructed the mutants Δ*popB*Δ*fliC* and Δ*popB*Δ*pilT* and quantified their engulfment by CFUs in THP-1 macrophages. Δ*popB*Δ*fliC* showed a pronounced defect in phagocytosis while Δ*popB*Δ*pilT* uptake was only mildly reduced, in alignment with the Tn-seq data (Fig. 1f). Next, we assessed the fitness of biofilm-associated genes. Mutations impairing Psl polysaccharide synthesis or lowering c-di-GMP levels (e.g., *wspR*) caused accumulation in both supernatant and intracellular environments, while mutations that elevate c-di-GMP and promote biofilm formation (e.g., *bifA*)[38] had the opposite effect, indicating general fitness effects during competition with macrophages (Fig. 1g,h). We next validated those effects on intracellular fitness with the mutants Δ*popB*Δ*pslBCD*, deficient in Psl secretion, and Δ*popB*Δ*bifA*, which displays a hyper-aggregation phenotype induced by high c-di-GMP levels. As observed in the Tn-seq data, intracellular CFUs showed enrichment of Δ*popB*Δ*pslBCD*, however no significant difference was observed for Δ*popB*Δ*bifA* (Fig. 1i). This inconsistency with the Tn-seq data may be due to the reduced sensitivity of CFU assays compared to Tn-seq and the hyper-aggregation phenotype of Δ*bifA*, which imposes technical challenges on CFU experiments.

Together, these findings implicate that *P. aeruginosa* can rapidly evade phagocytosis by loss of motility or induction of biofilm formation, providing a rationale for the selective pressure inactivating these systems during chronic infections. We next investigated the mechanisms by which motility systems influenced phagocytosis at the single cell level using high-resolution live microscopy.

### *P. aeruginosa* surface colonization facilitates phagocytosis

To map the contributions of bacterial motility in driving phagocytic processes, we turned to live microscopy to dynamically study bacteria-macrophage interactions. We used alveolar macrophage-like (AML) cells[39,40] as a model system to provide a relevant immune cell for modeling *P. aeruginosa* chronic infection of the airways. We first assessed bacterial uptake by AML cells with the control strain Δ*popB*. Macrophages were fluorescently labeled with CellTracker before inoculation with *P. aeruginosa* constitutively expressing the red fluorescent protein mScarlet. We found that upon exposure to Δ*popB*, macrophages initiated uptake within minutes. AML cells engulfed bacteria from planktonic and surface-associated populations via membrane protrusions (Fig. 2a).

**Figure 2:**
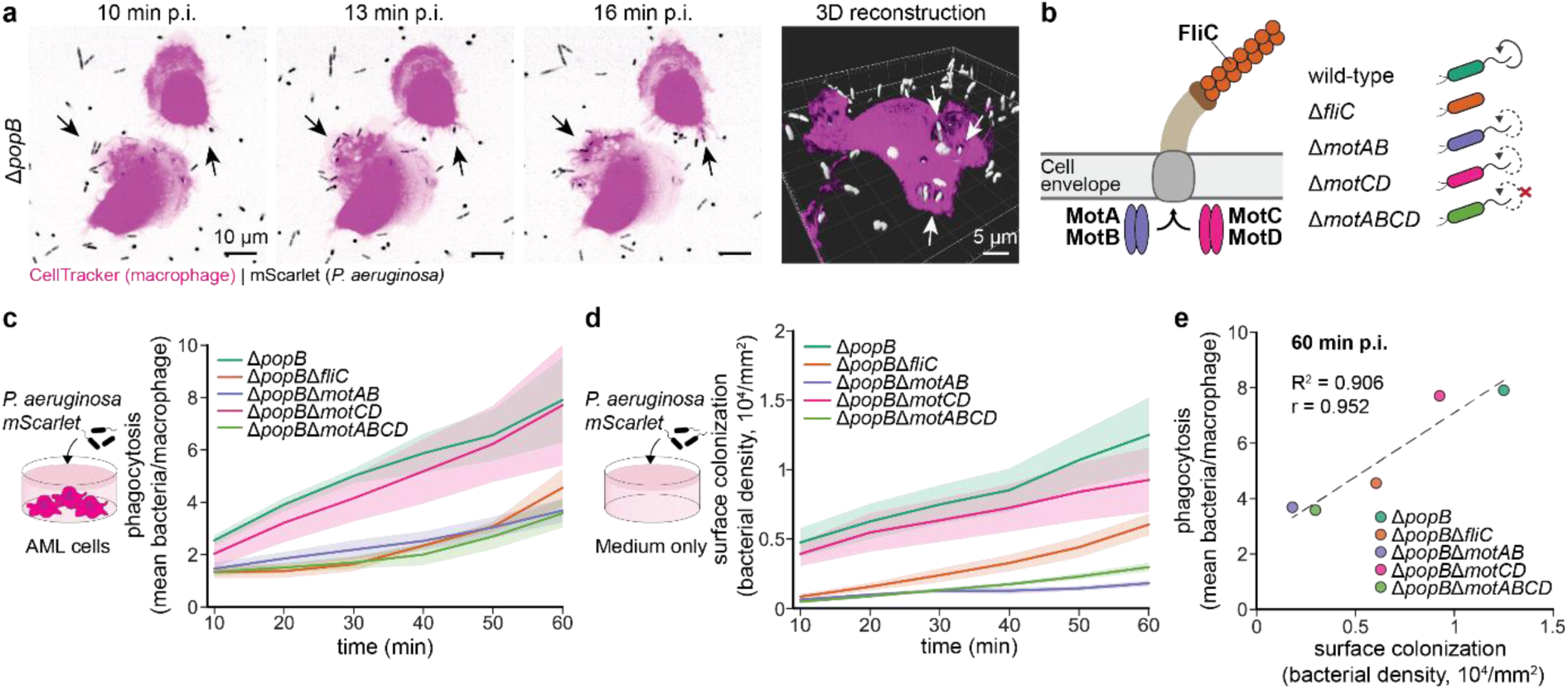
Surface colonization of swimming bacteria facilitates phagocytosis. **a**, Timelapse images of bacterial phagocytosis by AML cells at the surface (cropped image, first z-plane). AML cells were labelled with CellTracker and infected with the mScarlet-expressing PAO1 Δ*popB* mutant at MOI 100. On the right, a 3D reconstruction of a phagocytosing AML cell is shown. Arrows indicate engulfed bacteria. **b**, Depiction of flagellar composition and mutant phenotypes. The MotAB and MotCD stator complexes drive flagellar rotation, with differential recruitment depending on mechanical load. Loss of both complexes abolishes flagellar movement, whereas deletion of *fliC* prevents formation of the flagellum. **c**, Quantification of microscopy-based phagocytosis assay. AML cells were infected with different flagellar mutants at MOI 10 and timelapses were acquired for 1 h at a 10 min interval (4 z-positions, 2 µm step size). Cell segmentation was performed on the CellTracker and mScarlet signal for macrophages and bacteria, respectively. The mean number of intracellular bacteria per macrophage was calculated for each replicate and timepoint across the entire z-stack. Thick line, mean; shaded area standard deviation across biological replicates (n=3). **d**, In parallel to the phagocytosis timelapse, the same inoculum of bacteria was imaged in the absence of macrophages. Bacterial density at the surface (first z-plane) was quantified over time describing the dynamics of bacterial surface colonization. Thick line, mean; shaded area standard deviation across biological replicates (n=3). **e**, Correlation of phagocytosis and bacterial surface colonization at the end of the timelapse (60 min p.i.). Dashed line, linear fit; r, Pearson’s correlation coefficient (p=0.013); R^2^, coefficient of determination.

We quantified the uptake of flagellar mutants during the first hour of infection at MOI 10 by segmenting and computing the average number of intracellular bacteria per macrophage. Live-cell microscopy confirmed that bacterial uptake was strongly impaired for the mutant Δ*popB*Δ*fliC*. We next asked whether the flagellum or its function in swimming motility promoted phagocytosis. To answer this question, we relied on stator mutants impacting swimming motility but not flagellar expression. The mutant Δ*popB*Δ*motABCD* lacks both flagellar stator complexes which fully abolishes swimming motility. In contrast, Δ*popB*Δ*motCD* maintains wild-type swimming motility in low-viscosity environments while Δ*popB*Δ*motAB* slows down[41]. In comparison to Δ*popB*Δ*fliC*, bacterial uptake was similarly impaired for the mutants Δ*popB*Δ*motAB* and Δ*popB*Δ*motABCD* with swimming defects. Conversely, Δ*popB*Δ*motCD* was as efficiently phagocytosed as the control strain confirming intact swimming motility (Fig. 2b,c, S3). In summary, these results support previous findings that phagocytic evasion of *P. aeruginosa* is indeed driven by impaired swimming motility rather than loss of flagellum.

For phagocytosis to occur, bacteria must reach the local environments of surface-attached AML cells, which move at a slower pace compared to bacterial swimming (minutes versus seconds). We therefore hypothesize that defects in swimming motility limit *P. aeruginosa*’s ability to reach the surface, impeding macrophage encounters. To test this, we quantified bacterial accumulation on the substrate surface in the absence of macrophages. The non-flagellated Δ*popB*Δ*fliC* mutant showed a 2-fold decrease in attachment rate relative to Δ*popB* after 60 min of incubation. Mutations impacting motility while maintaining flagellar expression (Δ*popB*Δ*motAB* and Δ*popB*Δ*motABCD*) showed an even stronger defect (7-fold and 4-fold, respectively). In contrast, Δ*popB*Δ*motCD* surface colonization was only mildly reduced compared to Δ*popB* (Fig. 2d). Overall, we found a strong correlation between bacterial uptake and surface attachment (Fig. 2e). Together, our data supports a model where swimming motility facilitates bacteria-macrophage encounters improving phagocytic efficacy.

### Swimming motility promotes bacterial attachment to macrophages

Intrigued by the quantitative differences in surface colonization of flagellar mutants, we wondered whether swimming defects similarly modulate direct contacts between *P. aeruginosa* and macrophages. To explore this, we tracked bacterial binding events with AML cell membranes by timelapse microscopy at high temporal resolution, which allowed the quantification of contact durations and the stability of attachment (Fig. 3a,b).

**Figure 3:**
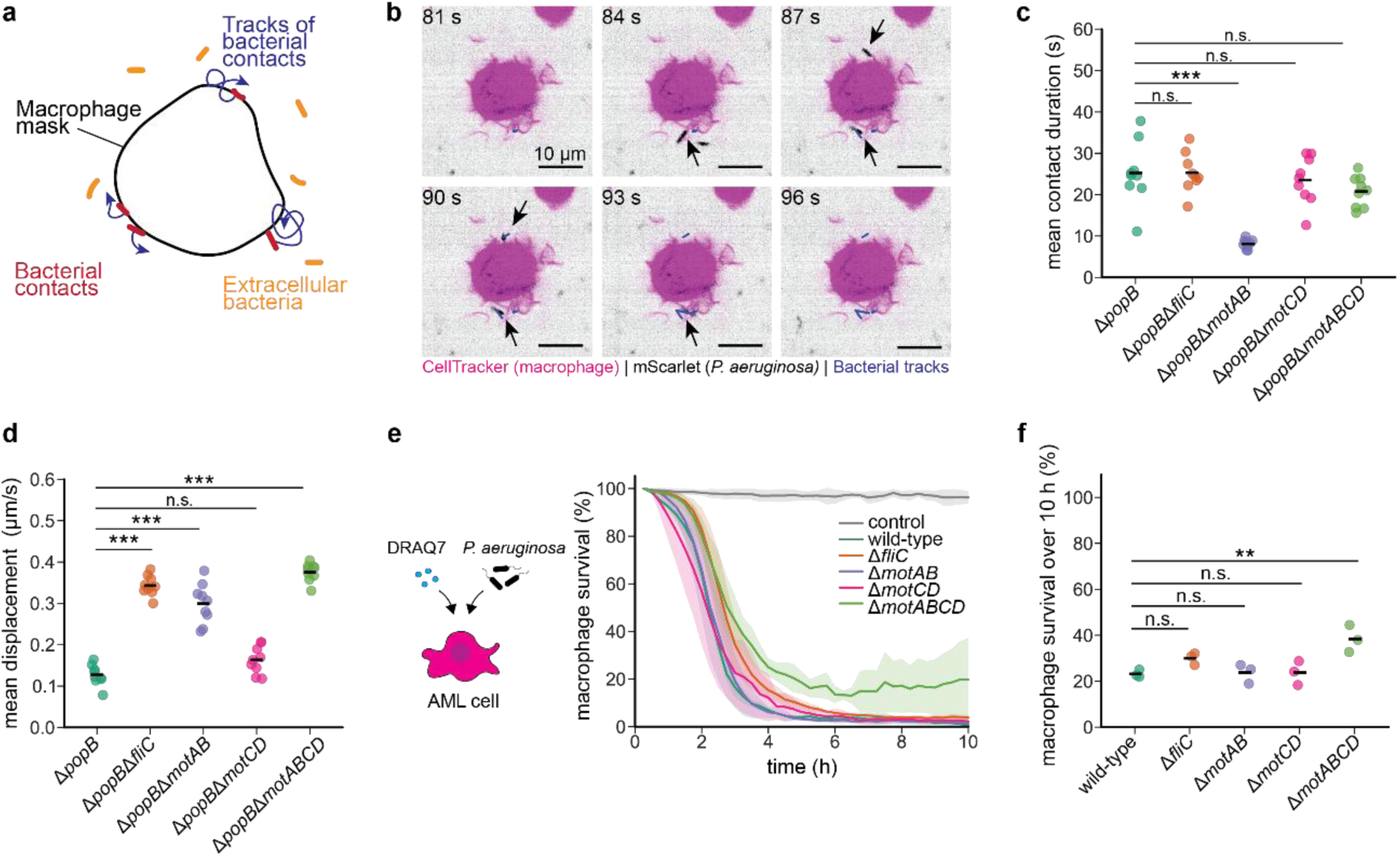
Attachment to macrophages is impaired in swimming mutants. **a**, Illustration describing the image analysis of bacteria-macrophage interactions. AML cells stained with CellTracker and mScarlet-expressing bacteria were recorded for 3 min at a 3 s interval (MOI 200, single z-plane above the surface). Cell segmentation of bacteria and macrophages was performed. Macrophage masks were enlarged to capture bacteria sitting on the macrophage surface. A filtering step was performed on bacterial masks overlapping with the macrophage masks which were defined as bacterial contacts while extracellular bacteria were excluded from the analysis. During contact with macrophages, bacteria were tracked providing information about contact durations and bacterial trajectories. **b**, Exemplary time series showing two bacteria getting into contact with a single macrophage and their tracks (cropped image, single z-plane). **c**, Bacterial contact durations were measured. Contacts that lasted for the entire duration of image acquisition were excluded since these were predominantly internalized bacteria. The mean contact duration for each replicate per mutant is depicted. Black line, mean; circles, technical replicates (from 3 independent experiments); statistical test, one-way ANOVA with Dunnett’s post-hoc. **d**, Bacterial displacement for each 3 s timeframe excluding gaps was calculated for all tracks of a replicate and the mean displacement was computed. Tracks spanning the whole imaging period were excluded. Black line, mean; circles, technical replicates (from 3 independent experiments); statistical test, one-way ANOVA with Dunnett’s post-hoc. **e**, Quantification of AML cell survival when infected with flagellar mutants in the more virulent PAO1 wild-type background (MOI 10). Thick line, mean; shaded area, standard deviation across biological replicates (n=3). **f**, Total survival over 10 h was calculated as the percentage of the bacterial strain AUC relative to the control AUC (uninfected cells). Black line, mean; circles, biological replicates; statistical test, one-way ANOVA with Dunnett’s post-hoc.

We hypothesized that reduced contact durations in swimming mutants may decrease the probability of bacterial engulfment which can take up to several minutes. We found that the average contact duration of Δ*popB*Δ*fliC*, Δ*popB*Δ*motABCD* and Δ*popB*Δ*motCD* was similar to Δ*popB*, while it was reduced for Δ*popB*Δ*motAB* (Fig. 3c). Strikingly, the distribution of contact durations showed that Δ*popB*Δ*motAB* had a particularly high prevalence (57 %) of very short contacts lasting only one imaging interval (3 s) relative to Δ*popB* (33 %) (Fig. S4).

Next, we looked at the mean displacement of tracks per imaging interval. A high average displacement suggests that bacterial attachment was weaker. We found that those mutants with swimming deficits (Δ*popB*Δ*fliC*, Δ*popB*Δ*motAB*, Δ*popB*Δ*motABCD*) were increasingly displaced during contact with macrophages compared to Δ*popB*. Furthermore, calculation of the mean square displacement during macrophage encounters revealed a diffusive and a modestly superdiffusive behavior of the immotile mutants Δ*popB*Δ*fliC* and Δ*popB*Δ*motABCD*, respectively. Δ*popB*Δ*motAB* which partially retains swimming motility showed a mild subdiffusive behavior but was largely constrained. In contrast, Δ*popB*Δ*motCD* presented a lower average displacement and was locally confined, similar to Δ*popB* (Fig. 3d, S5). For bacterial tracking, we decided to treat reoccurring contacts with the same macrophage as a single contact, thus explaining the longer contact durations of non-swimming mutants colliding multiple times with a macrophage’s membrane while diffusing (Movie S7, S10). In summary, shortened contact durations of Δ*popB*Δ*motAB* and the increased displacement of swimming mutants during contact further hinders efficient engulfment.

As contact between bacteria and macrophages is also required for injection of T3SS effectors, we wanted to know whether less stable interactions in swimming mutants could also reduce their cytotoxicity. To test this, we infected AML cells with swimming mutants in the PAO1 wild-type background. Compared to THP-1 macrophages, AML cells are more susceptible to T3SS-mediated killing by *P. aeruginosa* (Fig. S6a,b). Cytotoxicity assays in flagellar mutants revealed no significant reduction in killing except for Δ*motABCD*, which showed a minor delay (Fig. 3e,f). Hence, even transient and unstable contacts were sufficient for bacteria to engage their T3SS and rapidly eliminate macrophages. In the broader context of immune competition, these data suggest that loss of swimming motility provides a fitness advantage primarily under chronic, attenuated virulence conditions, allowing bacteria to avoid confrontation with immune cells. The scenario flips in acute strains with a functional T3SS: bacterial cytotoxicity outweighs susceptibility to phagocytosis, allowing bacteria to rapidly dominate over macrophages.

### Type IV pili regulate phagocytotic uptake of chronic strains

Based on the Tn-seq data, we next investigated the function of T4P in phagocytosis. T4P enable bacterial attachment and movement over surfaces by twitching motility, promoting exploration and biofilm formation. T4P also function in surface sensing, which induces bacterial virulence programs through cAMP signaling[42–44]. We first focused on the mutant Δ*popB*Δ*pilA*, which lacks the major pilin PilA, thereby abolishing T4P formation and twitching motility. Timelapse microscopy in AML cells revealed uptake of Δ*popB*Δ*pilA* to a lower extent than Δ*popB*. To decouple twitching motility from T4P production, we generated the Δ*popB*Δ*pilT* mutant deficient in T4P retraction motor PilT abolishing motility but causing hyper-piliation. We found that phagocytosis of the hyper-piliated, non-twitching mutant Δ*popB*Δ*pilT* was also reduced relative to Δ*popB* (Fig. 4a,b, S3). Given the role of T4P in surface adhesion, we next tested if these mutants displayed defects in surface colonization. Quantification of surface-associated bacteria in the absence of macrophages revealed only a mild reduction for Δ*popB*Δ*pilA* and no detectable change for Δ*popB*Δ*pilT* (Fig. 4c). These results suggest that loss of twitching motility rather than T4P expression may promote escape from phagocytosis. Yet, limited availability of Δ*popB*Δ*pilA* on the surface may contribute to its phagocytic escape.

**Figure 4:**
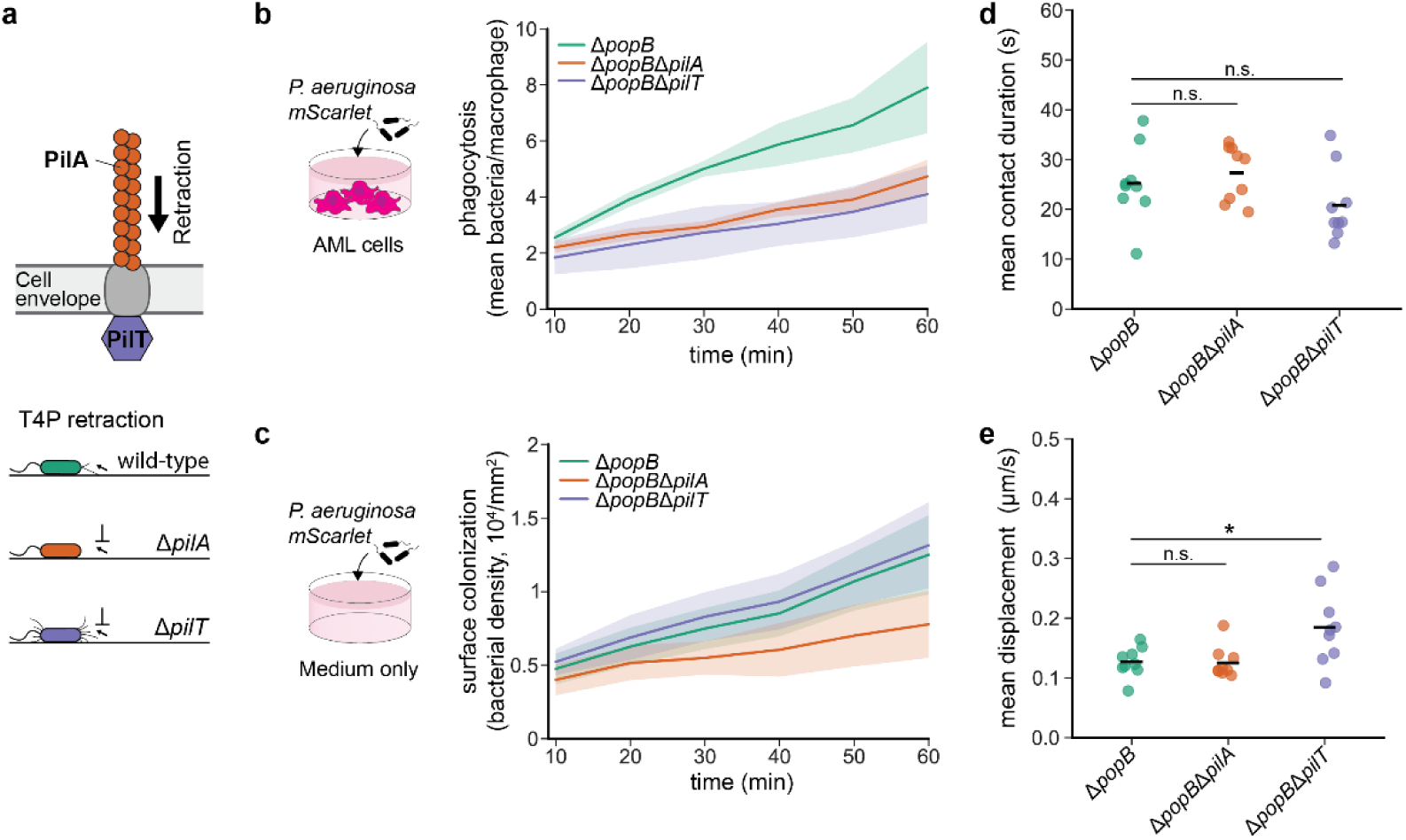
T4P biosynthesis and retraction modulate phagocytosis through distinct mechanisms. **a**, Illustration of the type IV pilus and mutant phenotypes. *pilA* deletion abolishes pilus formation, whereas loss of PilT prevents retraction resulting in hyper-piliation. **b**, Phagocytosis of bacterial mutants by AML cells at MOI 10 was quantified by macrophage and bacterial cell segmentation on timelapse imaging data (4 z-planes, 2 µm step size). Thick line, mean; shaded area, standard deviation across biological replicates (n=3). **c,** Surface colonization of the same bacterial inoculum used in the phagocytosis assay was monitored in the absence of macrophages. Thick line, mean; shaded area, standard deviation across biological replicates (n=3). **d**, Short timelapses of 3 min with a 3 s interval were acquired to assess dynamics of bacterial contacts with AML cells. The mean contact duration per replicate and mutant is shown excluding contacts that lasted the entire imaging period (internalized bacteria). Black line, mean; circles, technical replicates (from 3 independent experiments); statistical test, one-way ANOVA with Dunnett’s post-hoc. **e**, The mean displacement per time frame of 3 s (excluding gaps) was calculated across all tracks per replicate during bacteria-macrophage contact. Contacts spanning the entire timelapse were excluded. Black line, mean; circles, technical replicates (from 3 independent experiments); statistical test, one-way ANOVA with Dunnett’s post-hoc.

We next interrogated contact dynamics between *P. aeruginosa* and AML cells. No significant differences in bacterial contact durations were observed for Δ*popB*Δ*pilA* and Δ*popB*Δ*pilT* compared to the control (Fig. 4d, S4). However, Δ*popB*Δ*pilT* exhibited a significant increase in average displacement while bound to macrophage membranes, whereas Δ*popB*Δ*pilA* did not (Fig. 4e). Analysis of the mean square displacement over time further revealed that both Δ*popB*Δ*pilT* and Δ*popB*Δ*pilA* were locally confined (Fig. S5), indicating that the higher displacement of Δ*popB*Δ*pilT* reflects local fluctuations at the membrane rather than long-range diffusion.

Together, these results suggest that distinct mechanisms may cause impaired phagocytosis of Δ*popB*Δ*pilA* and Δ*popB*Δ*pilT*: Δ*popB*Δ*pilA* affects bacterial attachment and is less abundant on the surface reducing bacterial encounters with macrophages, whereas unstable binding of Δ*popB*Δ*pilT* at the membrane interferes with efficient uptake which is likely caused by hyper-piliation of this mutant. However, these partial defects in either surface colonization or macrophage attachment were rather mild. Hence, it is possible that other mechanisms come into play that do not directly modulate encounters between macrophages and bacteria but rather the phagocytic process itself. For instance, twitching motility may be sensed by macrophages amplifying immune responses.

### Piliation levels in non-swimming bacteria modulate phagocytic efficacy

Because twitching and swimming motility can be concomitantly abolished during chronic infection[45], we investigated the combined effect of impaired flagellar and T4P function. In the absence of a flagellum, hyper-piliation in the Δ*popB*Δ*fliC*Δ*pilT* mutant led to an even more severe phenotype which completely abolished phagocytosis by AML cells. In contrast, the non-piliated Δ*popB*Δ*fliC*Δ*pilA* mutant was internalized slightly more efficiently than the control (Fig. 5a, S3).

**Figure 5:**
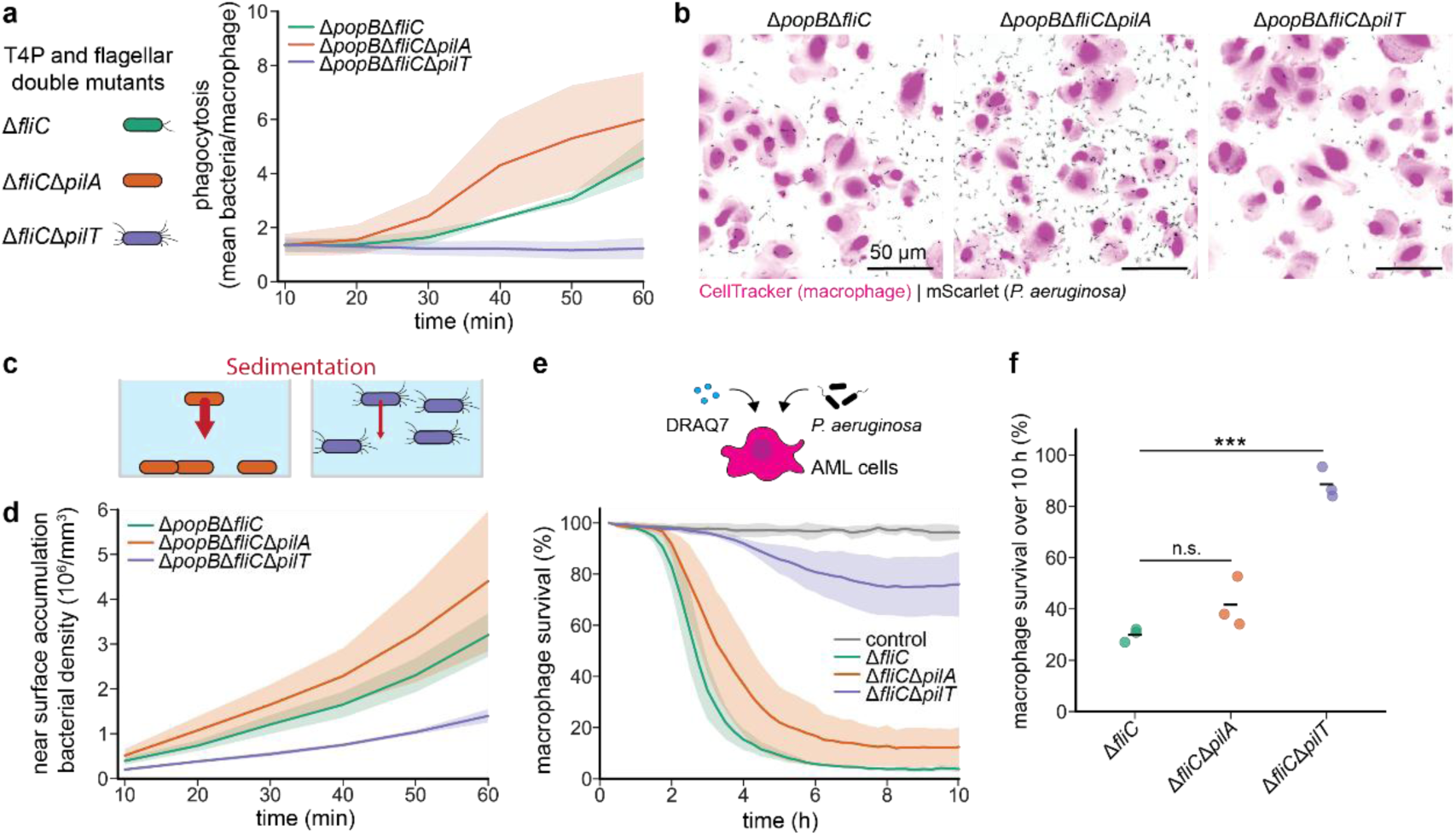
Hyper-piliation in non-flagellated *P. aeruginosa* exacerbates phagocytic defects and attenuates virulence. **a**, T4P mutants were constructed in the Δ*fliC* background to test the role of T4P functions in the absence of swimming. Results of the microscopy-based phagocytosis assay in AML cells are shown (MOI 10). Thick line, mean; shaded area, standard deviation across biological replicates (n=3). **b**, Representative maximum z-projections (cropped images, 4 z-planes, 2 µm step size) of the phagocytosis assay at 60 min p.i. showing the accumulation of bacteria near the surface and macrophages. **c**, Proposed model of accelerated sedimentation of the non-piliated (Δ*popB*Δ*fliC*Δ*pilA*) versus stalled sedimentation in the hyper-piliated mutant (Δ*popB*Δ*fliC*Δ*pilT*). **d**, Quantification of bacteria near the surface in the absence of macrophages measured across the z-stack (4 z-planes, 2 µm step size). The same inoculum as for the phagocytosis assay was used. Thick line, mean; shaded area, standard deviation across biological replicates (n=3). **e**, AML cell survival was quantified over time. Macrophages were infected with T4P mutants in the more virulent PAO1 wild-type background that were either flagellated or non-flagellated (MOI 10). Thick line, mean; shaded area, standard deviation across biological replicates (n=3). **f**, The total survival over 10 h is depicted for each bacterial strain representing the percentage of the AUC compared to the control (uninfected cells). Black line, mean; circles, biological replicates; statistical test, one-way ANOVA with Dunnett’s post-hoc.

In the absence of motility, bacteria must reach the surface and macrophages by sedimentation and Brownian motion. We suspect that hyper- and non-piliated bacteria show differences in sedimentation due to drag exerted by T4P, thereby impacting phagocytosis. We quantified bacteria in the vicinity of the surface without macrophages being present. Δ*popB*Δ*fliC*Δ*pilA* mutants showed an increased rate of accumulation near the surface. By contrast, the Δ*popB*Δ*fliC*Δ*pilT* mutant only rarely reached the surface (Fig. 5b,c,d). Thus, the impact of piliation on bacterial sedimentation modulates access to immune cells in liquid conditions.

We next assessed the competition of T4P mutants with macrophages in the highly virulent PAO1 wild-type background. Cytotoxicity assays showed rapid killing of AML cells by T4P mutants with intact flagellum (Fig. S6c,d). This was expected since both mutants were still able to make contact with macrophages facilitated by swimming motility. In contrast, sedimentation effects may influence bacterial virulence by preventing or accelerating macrophage encounters. Indeed, Δ*fliC*Δ*pilT* was strongly attenuated in virulence with 76 % of AML cells remaining viable at 10 h post-infection. Conversely, macrophage killing of Δ*fliC*Δ*pilA* was comparable to that of Δ*fliC* (Fig. 5e,f). This result suggests that the non-flagellated, hyper-piliated mutant stays afloat even over longer durations, limiting T3SS engagement at the macrophage surface. This phenotype is particularly intriguing as it may represent an alternative mechanism underlying attenuated virulence in chronic *P. aeruginosa* isolates, that does not require loss of T3SS.

Taken together, our findings support a model in which bacteria escape phagocytosis by avoiding proximity to macrophages through impaired swimming motility and by modulating T4P deployment. Having established these different immune evasion strategies, we next turned to clinical isolates to assess the relevance of our results.

### Clinical isolates recapitulate motility-dependent phagocytic evasion

Given the Tn-seq findings and mechanistic analyses identifying flagellar and T4P motility as key determinants of phagocytosis, we next asked whether these effects translate to clinically-relevant strains. From a global collection of *P. aeruginosa* human infection isolates[46], we selected isolates carrying mutations in regulators of flagellar and T4P biosynthesis[47,48]. Premature stop codons were prioritized as they are expected to abolish gene function. Among flagellar regulators, only medium-impact missense mutations in *fleQ* were detected, whereas multiple isolates carried a premature stop codon in either *pilS* or *pilR*, genes of the two-component system controlling T4P expression.

To assess the functional consequences of these mutations, we evaluated swimming motility by soft-agar assay and twitching motility by stab assay (Fig. 6a,b). *fleQ* mutations were expected to affect flagellar expression and potentially abolish swimming, whereas *pilS* or *pilR* loss-of-function should primarily interfere with twitching motility. Consistent with these expectations, the isolate PAE0140 carrying a *fleQ* missense mutation showed complete loss of swimming while retaining normal twitching motility. Conversely, the two isolates with premature stop codons in *pilR* (IPC_1633 and IPC_534) could not twitch and yet maintained swimming motility. Surprisingly, the isolate IPC_57 that carried missense mutations in *fleQ* and *pilR* and a premature stop codon towards the end of the gene *pilS* retained swimming and twitching motility (Fig. 6c, S7). Some of the clinical isolates showed reduced expansion in swimming and twitching assays compared to the positive controls in the PAO1 lab strain background, which was likely due to slower bacterial growth (Fig. S8).

**Figure 6:**
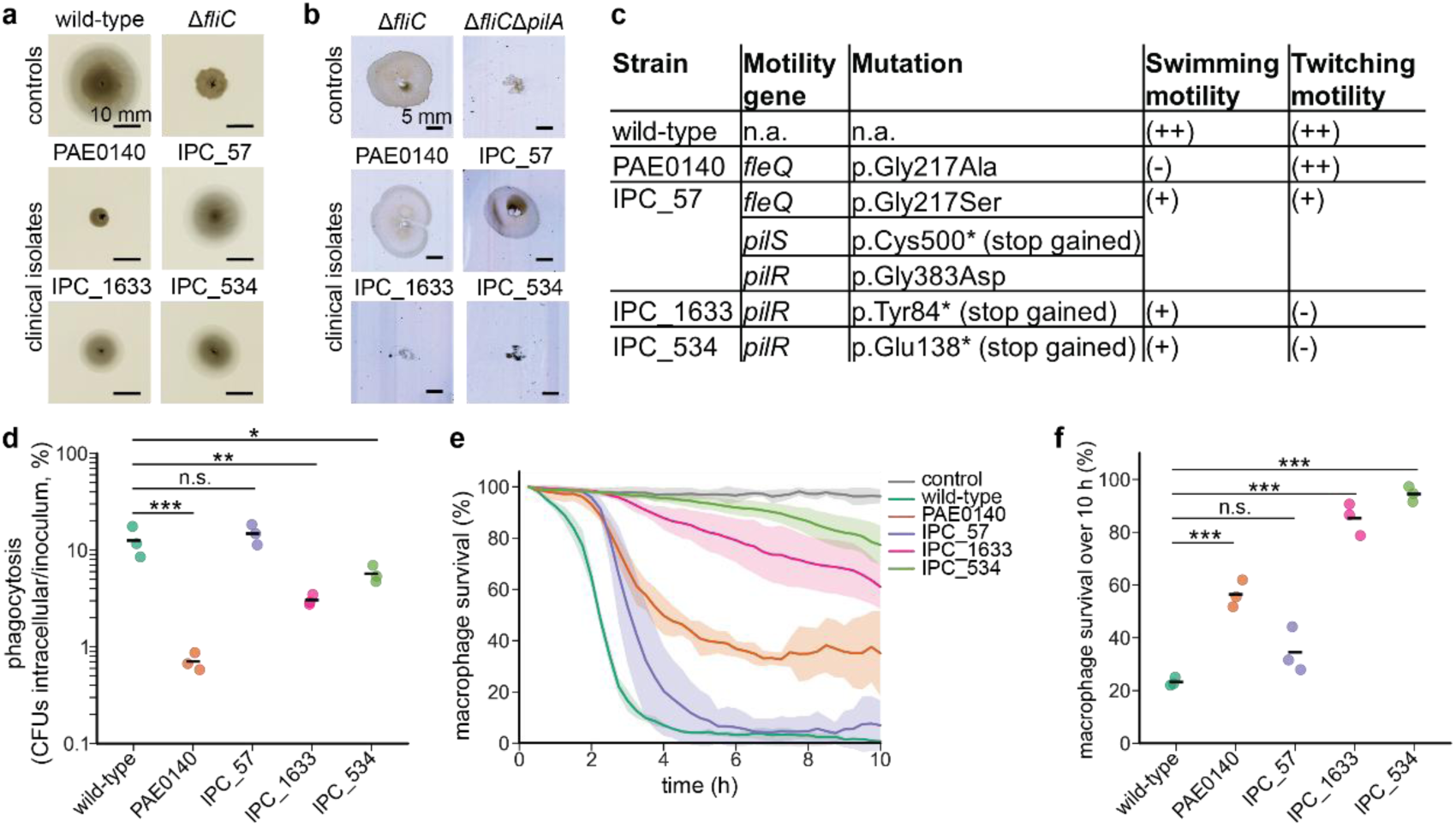
Phagocytic evasion of clinical isolates correlates with their swimming and twitching motility phenotypes. **a**, Results of the swimming plate assay in *P. aeruginosa* clinical isolates with PAO1 wild-type as a positive control and Δ*fliC* as a negative control. **b**, Twitching motility of clinical isolates was determined by stab assay using PAO1 Δ*fliC* as a positive and Δ*fliC*Δ*pilA* as a negative control. **c**, Table summarizing the results of the swimming and twitching motility assays. (++) indicates near-wild-type twitching or swimming motility, whereas mutants marked with (+) display reduced motility. (-) signifies no motility was detected. Additionally, information about clinical isolate mutations in T4P and flagellum-associated genes is provided. **d**, CFU-based phagocytosis assay in AML cells measuring uptake of clinical isolates compared to PAO1 wild-type (MOI 10). Intracellular bacteria were quantified 1 h p.i. without gentamycin selection. Black line, mean; circles, biological replicates (n=3); statistical test, one-way ANOVA with Dunnett’s post-hoc. **e**, Survival of AML cells is shown over time for each clinical isolate and PAO1 wild-type, with uninfected macrophages serving as a control. Thick line, mean; shaded area, standard deviation across biological replicates (n=3). **f**, Total survival over the entire infection period of 10 h is presented with the percentage comparing the AUC to the control (uninfected cells). Black line, mean; circles, biological replicates (n=3); statistical test, one-way ANOVA with Dunnett’s post-hoc.

We next tested whether motility phenotypes correlated with bacterial internalization. To this end, we quantified phagocytosis by CFU recovery from AML cell lysates after 1 h infection at MOI 10. Isolate PAE0140 showed markedly reduced uptake in line with its swimming motility defect, while the isolates IPC_1633 and IPC_534 displayed intermediate phagocytic defects as expected for non-twitching mutants. In contrast, phagocytosis of IPC_57 was comparable to our lab strain PAO1 wild-type (Fig. 6d). This was expected as isolate IPC_57 did not present any major motility defects.

Because earlier experiments in laboratory strains used the less virulent Δ*popB* background, we considered whether differences in cytotoxicity between clinical isolates might influence their uptake. If cytotoxicity had an impact on phagocytosis, we would expect a reduced recovery of intracellular bacteria in highly virulent isolates due to macrophage death. On the contrary, killing assays in AML cells revealed an opposite trend where isolates with reduced phagocytosis (PAE0140, IPC_1633, IPC_534) also exhibited attenuated cytotoxicity, while IPC_57 presented near-wild-type virulence (Fig. 6e,f). Thus, phagocytic evasion of clinical isolates could not be attributed to T3SS-mediated macrophage killing, supporting that motility is the primary driver of phagocytic differences in these clinical isolates. Together, our data demonstrate a correlation between bacterial motility and uptake by macrophages in clinically relevant *P. aeruginosa* strains.

## Discussion

Our functional genomics screen in an attenuated *P. aeruginosa* background showed a strong selection for motility mutants. Swimming motility in particular has long been recognized as an important determinant for phagocytosis. By employing live-cell imaging of bacteria-macrophage interactions, we identified the biophysical mechanisms by which motility impacts phagocytosis. Swimming bacteria move much faster than macrophages, increasing the frequency of encounters. Engulfment occurs either upon direct contact of bacteria with the macrophage membrane or by collecting bacteria from the surface. We showed that loss of swimming motility impairs phagocytosis by interfering with surface colonization and attachment to macrophages. Moreover, we identified different contributions of the flagellar stator complexes MotAB and MotCD. While Δ*motAB* was poorly internalized and showed defects in surface colonization and macrophage binding, Δ*motCD* behaved like the wild-type. On the one hand, swimming motility is differently affected in these two mutants with Δ*motCD* retaining wild-type swimming motility in low viscosity environments such as the cell culture medium used here, whereas Δ*motAB* swimming motility is impaired[41]. On the other hand, only the stator complex MotAB is required for flagellum-mediated surface sensing and adaptation during initial contact[49]. Thus, our results support a model where effective phagocytic uptake requires a functional flagellum. Thereby, swimming motility increases the probability of contacts, while the flagellum simultaneously participates in surface sensing enabling prolonged binding to macrophages and the substrate surface. A reduction in bacteria-macrophage interactions may also explain why previous studies observed altered inflammatory responses to non-swimming mutants with macrophages presenting decreased PI3K/Akt and inflammasome signaling[50,51].

We also uncovered a previously underappreciated role of T4P in phagocytosis. Both non-piliated and hyper-piliated mutants evaded engulfment by AML cells. Reduced uptake of non-piliated mutants reflected mild defects in surface colonization, while the hyper-piliated mutant was impaired in establishing stable contacts with macrophages. Several studies have identified an important function of T4P in attachment to biotic and abiotic surfaces, however binding profiles vary significantly with different surface composition such as distinct motifs present on mammalian cells[52–54]. Furthermore, T4P mediate surface adaptation and biofilm formation through mechanosensory pathways enforcing bacterial adhesion over extended periods[42,55]. We found that hyper-piliation tends to inhibit bacterial adhesion to macrophages, a behavior opposite to attachment on abiotic surfaces. Hence, we propose a model where hyper-piliation may sterically hinder engagement of bacteria with macrophage receptors and binding motifs that are specific to other bacterial ligands than T4P. Another possibility is that T4P retraction may mechanically stimulate macrophages, which would explain the reduced uptake of both non-twitching mutants. There is precedence of mechanical activation of host cells in *Neisseria gonorrhoeae,* which amplifies infection-associated gene expression in epithelial cells through T4P retraction[56].

In the absence of swimming motility, loss of pili increased phagocytosis, while hyper-piliation had the opposite effect and strongly diminished engulfment. The large number of T4P in hyper-piliated, non-swimming mutants increase a cell’s hydrodynamic drag, slowing down sedimentation and thereby dramatically decreasing access to surface-attached predators[57]. Strikingly, in acute conditions, Δ*fliC*Δ*pilT* induced a virulence deficit quantitatively identical to a T3SS loss of function mutation like Δ*popB*. In line with these results, murine and *Caenorhabditis elegans* infection models exhibit improved survival with hyper-piliated *P. aeruginosa*, consistent with impaired toxin delivery due to insufficient host cell contact[58,59]. Hyper-piliation has been reported in CF small-colony variants[60] where it facilitates auto-aggregation and biofilm formation[61]. Yet, Tn-seq studies identified advantages only for non-piliated, but not hyper-piliated mutants during *P. aeruginosa* colonization of the mouse lung and gastrointestinal tract[62,63]. These discrepancies highlight the context dependence of T4P phenotypes with distinct effects on host colonization and immune evasion, which are further modulated by other traits including swimming motility. Analysis of clinical isolates in our infection model however reinforced our findings: impaired swimming or twitching motility coincided with reduced bacterial internalization by macrophages.

The rise of antimicrobial resistance poses significant challenges to treatment of bacterial infections. A better understanding of evasion strategies, including how motility influences bacterial interactions with immune cells, can inform approaches to enhance the body’s natural defenses against pathogens such as *P. aeruginosa*. Future work should examine the mechanisms by which T4P activity modulates macrophage function. Importantly, exploring immune functions in physiologically realistic microenvironments, such as mucosal airway models[64–66], will be key to understanding how bacterial behaviors shape host-pathogen interactions.

## Methods

### Bacterial culture

Bacteria were cultured in LB medium (Roth) in a shaking incubator at 37 °C and 225 rpm. LB agar plates were prepared by adding 1.5 % agar (Fisher scientific) to LB medium. For *E. coli* selection, 100 µg/ml ampicillin (Huberlab), 100 µg/ml carbenicillin (Fisher scientific) or 10 µg/ml gentamycin (Biochemica) were used. For *P. aeruginosa* selection, 300 µg/ml carbenicillin, 60 µg/ml tetracycline (Huberlab), 10 µg/ml chloramphenicol (Sigma-Aldrich) or 60 µg/ml gentamycin was added to the medium. For growth curves, bacteria in exponential phase were diluted to an OD_600_ of 0.01 in 200 µl LB medium. The optical density was recorded every 10 min during 20 h of bacterial growth at 37 °C using the Spectrostar plate reader (BMG Labtech). The plate was shaken at 300 rpm for 10 s before each measurement.

### Bacterial strains and vector construction

All strains and plasmids used in this study are listed in Table S1 and S2 in the supplementary information. Clinical isolates of *P. aeruginosa* were kindly provided by the Floto laboratory[46], selecting strains with mutations in the genes *fleQ*, *pilR* and *pilS*. *P. aeruginosa* PAO1 ATCC 15692 (American Type Culture Collection) was used for generation of all mutants. *E. coli* strains DH5α and S17.1 were used for vector construction and conjugation, respectively. Strains were constructed by two-step allelic exchange[67] using the suicide vectors pEX18_Amp_ or pEX18_Gent_ as previously described[36]. For genomic deletions, homology arms encompassing 500-1,000 bp upstream and downstream of the target gene were amplified by PCR and inserted into the suicide vector by Gibson assembly[68]. Following conjugation, marker-less deletions were confirmed by PCR and sequencing. Fluorescent strains for microscopy-based assays were generated by mini-Tn7 insertion[69] of mScarlet under the control of the *tet* promotor into the *att*Tn7 site as previously described[64].

### Mammalian cell culture

All mammalian cells were kept at 5 % CO_2_ and 37 °C during culture and live microscopy assays. THP-1 monocytes (ATCC TIB-202) were cultured in RPMI 1640 medium supplemented with 10 % FBS and 1X GlutaMAX (all from Thermo Fisher). Cells were split every 3 to 4 days. Respectively, 1 million, 200,000 or 20,000 THP-1 monocytes per well were seeded in a 6-well plate (Corning) for Transposon sequencing, 24-well (Corning) for CFU experiments or in a 96-well PhenoPlate (Revvity) for cytotoxicity assays. 100 nM phorbol 12-myristate 13-acetate (Adipogen) were added for differentiation to macrophages. Assays were performed 3 days after the start of differentiation.

Peripheral mononuclear cells (PBMCs) were isolated from human blood buffy coats (Transfusion Interregionale CRS). Buffy coats were diluted in MACS buffer (Miltenyi) at a ratio of 1:3 and 35 ml were layered on top of 15 ml Ficoll (Cytiva) in a 50 ml Falcon tube. After centrifugation at 400 g for 40 min in a swinging-bucket rotor without brake, the PBMC layer was carefully aspirated. PBMCs were washed in MACS buffer with centrifugation steps at 300 g and 200 g to remove platelets. Lastly, PBMCs were filtered through a 40 µm cell strainer (pluriSelect) and stored as aliquots in FBS supplemented with 10 % DMSO (Fisher scientific) in a liquid nitrogen tank.

For alveolar macrophage-like (AML) cell generation, PBMCs were thawed and, after a washing step in RPMI 1640, cells were labeled with human CD14 MicroBeads UltraPure (Miltenyi, 130-118-906). Monocytes were isolated following manufacturer’s instructions using LS columns and the MidiMACS separator (Miltenyi). Monocytes were then differentiated to AMLs in a 60 ml Teflon jar (Savillex) as previously described[70]. Briefly, 14 million monocytes were cultured in 7 ml RPMI 1640 supplemented with 10 % human serum (Sigma-Aldrich). 100 µg/ml Infasurf (Onybiotech), 10 ng/ml GM-CSF, 5 ng/ml TGF-β1 and 5 ng/ml IL-10 (Preprotech; 300-03-20UG, 100-21C-10UG, 200-10-10UG) were added to the medium on day 0, 2, 4 of differentiation and on day 6 when seeding AML cells. For seeding, AML cells were placed on ice for 30 min in the Teflon well. Cells were harvested and the Teflon well was washed thrice with ice-cold RPMI 1640. AML cells were seeded at 100,000 cells per well in 8 Well^high^ ibiTreat µ-slides (Ibidi) for microscopy-based phagocytosis and bacterial contact assays, at 30,000 cells per well in a 96-well PhenoPlate for the cytotoxicity assay, and at 200,000 cells per well in 24-well plates to assess phagocytosis by CFUs. Assays were performed on day 1 or 2 after seeding.

THP-1 macrophage infections for the Tn-seq screen were performed in RPMI 1640 supplemented with 10 % FBS which may contain low levels of active complement. Mechanistic experiments with AML cells were conducted in RPMI 1640 supplemented with 10 % heat-inactivated human serum that was additionally filtered at 100 kDa to avoid confounding effects of complement- and immunoglobin-mediated opsonization.

### Gentamycin exclusion assay

To isolate intracellular bacteria from macrophages, gentamycin exclusion assay was performed. Shortly, bacterial cultures in expansion phase were pelleted and resuspended in PBS. The optical density 600 (OD_600_) was adjusted to 1, and 5 µl of bacteria were added per 200,000 THP-1 macrophages in 500 µl cell culture medium (RPMI 1640 supplemented with 10 % FBS) to match a MOI of 10. After 1 h incubation at 37 °C, macrophages were washed with PBS and incubated for another hour with 60 µg/ml gentamycin in cell culture medium. Finally, 3 washes with PBS were performed and macrophages were lysed with 0.1 % Triton (VWR) in PBS for 10 min. Bacteria were collected in lysis buffer and further processed for the respective assay.

### Tn-seq assay

#### Transposon library construction

The Tn5 transposon T8 (IS*lacZhah*-tc) was conjugated into the *P. aeruginosa* PAO1 Δ*popB* strain as previously described[65,71,72]. Due to low conjugation efficiency, volumes had to be upscaled. Shortly, 100 ml overnight cultures of the *E. coli* donor strain SM10λ*pir* comprising the plasmid pIT2 was prepared in 100 µg/ml carbenicillin. The culture was pelleted, washed with LB and adjusted to an OD_600_ of roughly 160. 16 cultures of 5 ml each with the Δ*popB* strain were grown overnight at 42 °C without shaking. The culture was pelleted and the OD_600_ was adjusted to a ratio of approximately 1:3 to the donor strain. 2 ml of each bacterial culture were mixed together and 80 puddles of each 50 µl were spotted on LB plates and incubated for 2.5 h at 37 °C. Puddles were scraped off with an inoculation loop and pooled together in 16 ml LB medium. 200 µl aliquots were plated per LB agar plate supplemented with 60 µg/ml tetracycline and 10 µg/ml chloramphenicol and incubated for 24 h at 37 °C. Roughly 450,000 single colonies were harvested in 15 % glycerol (Sigma-Aldrich) in LB medium and were stored in 1 ml aliquots at -80 °C. The final stock had an OD_600_ of 10.

#### Tn-seq experimental design

To assess selection of *P. aeruginosa* during phagocytosis, the gentamycin protection assay was performed in the presence of the Transposon library and THP-1 macrophages. To screen for uptake of bacterial mutants by macrophages, both the supernatant and intracellular bacteria (lysate) were collected. Briefly, the transposon library was thawed in the morning, diluted to an OD_600_ of 0.1 and grown for 3.5 h to an OD_600_ of 0.48. Bacteria were then pelleted and adjusted to an OD_600_ of 1. 25 µl of the Transposon library culture were added per well of 1 million THP-1 macrophages in 1 ml culture medium. 1 h post-infection, the supernatant from 3 wells was pooled followed by a regrowth step. At 2 h post-infection, macrophages were lysed and intracellular bacteria from 3 wells were pooled together, again followed by a regrowth step. In parallel, the inoculum of the transposon library used for infection of macrophages was also regrown as a control. To match the number of generations between samples, all cultures were regrown to a similar OD_600_ between 0.4 to 0.45. They were then pelleted and frozen at -20 °C for further processing. The freshly thawed transposon library and the inoculum used for infection without regrowth were collected as additional controls. Bacteria were cultured throughout the entire experiment in macrophage culture medium (RPMI 1640 supplemented with 10 % FBS) including growth steps to avoid a selection bias due to changes in media composition.

#### Transposon library preparation

Library preparation was performed as previously described[65]. gDNA was isolated from bacterial pellets using the QIAamp DNA Mini Kit (Qiagen). Library preparation and sequencing was performed by Lausanne Genomic Technologies Facility (University of Lausanne). Genomic DNA (500 ng) was first sheared with a Covaris S220 using 400 bp insert settings (50 µL in microTUBES with AFA fiber, Peak incident power: 175, Duty factor: 5 %, Cycles per burst: 200, time: 70 s). Libraries were prepared with the xGen DNA MC UNI Library Prep Kit (IDT, protocol version v2) using xGen UDI-UMI adapters (IDT, 15 µM stock). With these adapters, P5 and P7 sequence are inverted compared to Illumina adapters, allowing transposon sequencing directly from read 1 (P5 side) in a single-end run (see below).

The purified ligated product was PCR-amplified with a primer specific for the Illumina P7 sequence (CAAGCAGAAGACGGCATACGA) and a second one specific for the transposon sequence (CGACGTTGTAAAACGACCACGT) carrying a 5’-biotin. PCR was performed with the KAPA HiFi HotStart ReadyMix kit (Roche). Cycling conditions were 98 °C for 45 s, followed by 10 cycles of 98 °C for 15 s, 60 °C for 30 s, and 72 °C for 30 s, and a final extension of 1 min at 72°C. The library was purified with SPRI beads at a 1X ratio.

The PCR product was captured with pre-washed Dynabeads MyOne Streptavidin T1 (Thermo Fisher). Binding & Wash (B&W) Buffer 2X composition is 10 mM Tris-HCl pH 7.5, 1 mM EDTA, 2 M NaCl. At least 1 ml of 1X B&W Buffer was added and mixed with 25 µl of Dynabeads. After 1 min on a magnet, the supernatant was removed and Dynabeads washed with the same volume of 1X B&W Buffer. This wash was repeated a second time. After removal of the supernatant on the magnet, the Dynabeads were resuspended with 50 µl of 2X B&W Buffer.

50 µl of library was mixed with the washed Dynabeads followed by incubation at RT on a rotator for 30 min. After 2 min on a magnet and discard of the supernatant, a wash was done with 100 µl of 1X B&W Buffer. Two additional washes were done before the final elution in 40 µl H2O. Half of the washed capture was used for the nested PCR with the Illumina P7 sequence (see sequence above) and a tailed primer made of the Illumina P5 sequence (red), the TruSeq read 1 primer binding site (black), and a transposon specific binding sequence (orange) AATGATACGGCGACCACCGAGATCTACACTCTTTCCCTACACGACGCTCTTCCGATCTCCAGGACGCTACTTGTGTAT. This nested PCR amplification was performed with the KAPA HiFi HotStart ReadyMix kit with the same cycling conditions as above but 8 cycles. The final library was purified with SPRI beads at a 0.8X ratio. It was quantified with a fluorimetric method (Qubit, Thermo Fisher) and its size pattern analyzed with a fragment analyzer (Agilent).

Sequencing was performed on a NovaSeq 6000 (Illumina) on a lane of an SP flow cell for a 100 cycles single end sequencing run. Clustering was performed with 1.2 nM library spiked with PhiX (Illumina). Sequencing data were demultiplexed using the bcl2fastq conversion software (version 2.20, Illumina).

#### Tn-seq data processing

“Wig” files after deduplication removing PCR duplicates by unique molecular identifiers (UMIs) that contained only primary reads were used for further data processing in TRANSIT[73] (v3.3.3). First, the quality of the data was assessed in TRANSIT showing no unexpected behavior (skewness of data within normal range). The Tn5 “resampling” method in TRANSIT was used to perform pairwise comparisons between conditions. The regrowth culture of the inoculum served as a reference and was compared to the samples (1) supernatant regrowth and (2) intracellular fraction regrowth. The recommended normalization method trimmed total reads (TTR) together with the default parameters was applied (samples=10,000, pseudocount=0, adaptive = True, histogram = False, include zeros = True, site-restricted=True, winsorize = False). We did not have sequencing replicates. However, each sample was pooled from several wells of macrophages accounting for biological variability. Statistical significance of fold-changes between conditions was determined by the adjusted p-value (applying the Benjamin Hochberg method) with a cut-off at p<0.05.

### Colony forming units

Intracellular bacteria were isolated from THP-1 macrophages by gentamycin exclusion assay described earlier. For CFUs in AML cells, selection with gentamycin was not possible due to variable resistance profiles of clinical isolates. AML cells were therefore directly lysed at 1 h post-infection after several washing step with PBS. Macrophage lysates and bacterial inoculum were diluted in PBS, and plated on LB plates. The plated dilutions were subsequently grown at 30 °C over night and bacterial colonies were quantified. For THP-1 cells, 3 to 4 technical replicates were prepared per biological replicate. Due to limited availability of AML cells isolated and differentiated from blood samples, only a single technical replicate was performed per biological replicate.

Phagocytosis was calculated as followed:

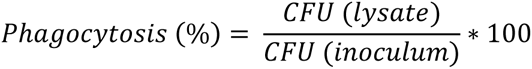

### Cytotoxicity assay

AML cells were stained for 30 min with 1 µM CellTracker green (Thermo Fisher) in plain RPMI 1640 prior to the experiment. Bacterial cultures in exponential phase were pelleted and resuspended in PBS. The OD_600_ was adjusted to 1. Per well of 30,000 AML cells in a 96-well plate, 100 µl RPMI 1640 medium supplemented with 10 % human serum (heat-inactivated, 100 kDa-filtered) and 3 µM DRAQ7 dye (Thermo Fisher) premixed with 0.75 µl culture with an OD_600_ of 1 (MOI 10) was added. Per experiment, 3 technical replicates were prepared for each mutant. Timelapse microscopy was acquired at the Operetta (PerkinElmer) microscope with a 20X water immersion objective (NA=1.0, binning 2) taking images at 15 min intervals in a single plane.

### Microscopy-based phagocytosis, surface colonization and contact assays

AML cells were stained for 30 min with 1 µM CellTracker green prior to the experiment. Bacterial cultures in exponential phase were resuspended in PBS and adjusted to an OD_600_ of 1. For phagocytosis assays, 200 µl RPMI 1640 supplemented with 10 % human serum (heat-inactivated, 100 kDa-filtered) premixed with 2.5 µl OD1 bacterial culture (MOI 10) were added to 100,000 AML cells in an Ibidi 8-well slide. In parallel, the same cell culture medium premixed with bacteria was added to empty wells without macrophages to assess bacterial surface colonization. All imaging data was acquired at a Nikon Ti2 confocal spinning disk microscope. Z-stacks were taken at 3 positions per well at an interval of 10 min during the first hour of infection using a 40X air objective (NA=0.95). For macrophage-bacteria contact assays, AML cells were instead infected at MOI 200. Only a single z-plane above the surface was acquired with a 60X water-immersion objective (NA=1.2) at an interval of 3 s for 3 min within the first 20 min of infection. The bacterial channel was imaged for all timepoints while for the macrophage channel a single image was taken at t=0 s.

### Swimming plate assay

LB medium with 0.3 % agar was prepared freshly (25 ml per 9 cm petri-dish). Bacterial overnight cultures were adjusted to an OD_600_ of 1. After letting the LB soft agar plates cool down for 30 min, 2 µl of bacterial culture were pipetted in the center of the agar layer. Plates were incubated for 24 h at 30 °C. In total 4 replicates were collected on 2 independent days. Data was recorded (Epson Perfection V850 Pro, Silverfast program) and the mean swimming diameter was measured in FIJI (version 1.54p).

### Twitching stab assay

Plates for stab assays were poured 24 h prior to experiments by preparing 0.5 % agarose, 5 g/l tryptone (Roth), 2.5 g/l NaCl (Fisher scientific) in bi-distilled water (33 ml per 9 cm petri-dish). After drying plates for 28 min, they were stored in the fridge. Bacteria were directly taken from a freshly streaked plate using a 10 µl pipet tip, stabbing vertically through the agar until reaching the bottom. Plates were incubated for 20 h at 37 °C, after which the agar was gently removed and twitching motility at the bottom of the plate was recorded (Epson Perfection V850 Pro, Silverfast program). Mean twitching diameters were measured in FIJI.

### Imaging data analysis

Preprocessing of image files including conversion from “nd2” to “tif” format and splitting of images into single channels, timepoints and z-planes was done in FIJI.

#### Phagocytosis

For the phagocytosis assay, each z-plane was analyzed individually. Macrophages were segmented with Cellpose[74,75] (version 3.0.10) using the cyto3 model (diameter 120, flow threshold 0.5, cell probability threshold 0). Bacteria were segmented with Omnipose[76] (version 1.0.7.dev77+g09ecee5) using the bact_fluor_omni model (flow threshold 0.2, mask threshold -3). To quantify intracellular bacteria, the macrophage masks were eroded by a disk of 10 pixels radius corresponding to 1.6 µm (scikit-image[77], skimage.filter.rank.minimum) to exclude bacteria attached to macrophages. Bacterial masks were multiplied with the binary image of the eroded macrophage mask and the number of remaining bacterial masks was quantified to calculate the number of intracellular bacteria. Lastly, the number of intracellular bacteria across all z planes was normalized by the number of macrophages to get an average of bacteria per macrophage. To quantify bacteria at the surface in the absence of macrophages, only bacterial masks in the first z plane were quantified. Both segmentation and quantification were performed in Python (version 3.9.21).

#### Bacterial contacts

For the macrophage-bacteria contact analysis, segmentation of bacteria was performed with Omnipose (bact_fluor_omni model, flow threshold 0.3, mask threshold -2) and segmentation of macrophages was done with Cellpose (cyto3 model, diameter 250, flow threshold 0.45, cell probability threshold 0). The macrophage mask was dilated by 10 pixels (1.1 µm) to include bacteria touching only the surface of the macrophage. Next, bacterial masks, macrophage masks and the original bacterial fluorescence images were merged for each timelapse and imported into TrackMate[78,79] in FIJI. Tracking was performed only on those bacterial masks which were overlapping with the macrophage masks (label image detector, spot filtering on mean intensity of macrophage mask). The simple LAP tracker was applied (maximum gap 3 µm, linking distance 3 µm, maximum frame gap 2). Subsequently, tracks were manually inspected and corrected by linking interrupted tracks and removing low quality tracks (out-of-focus bacteria). Data analysis on the exported track, spot and edge tables was performed in Python. Tracks that spanned the entire imaging period from 0 s to 180 s were excluded in the data analysis, since the majority of these tracks represented internalized bacteria. Contact durations were based on the track durations directly exported from TrackMate. For the distribution of contact durations all tracks from all replicates per mutant were pooled together. Additionally, the average contact duration was calculated for each replicate and mutant. To compute the mean displacement during bacterial contact, edges representing gaps were removed from the data set. Then, the mean displacement was calculated for each edge (representing the 3 s imaging time interval) and the average was determined from all track edges per replicate. To calculate the mean square displacement over time, the tracks of all replicates per mutant were pooled. All possible timepoint combinations per track were used to compute the mean square displacement for each lag time, while excluding timepoints representing a gap in the track. The lag time interval equalled the imaging interval of 3 s. Since tracks had different lengths, the number of available tracks decreased with increasing lag time. Thus, different cut-offs for linear curve fitting were chosen based on data availability (flagellar mutants, cut-off at 45 s; T4P mutants and control, cut-off at 90 s).

#### Macrophage survival

For the cytotoxicity assay, flatfield correction was done in the Harmony software (PerkinElmer) followed by data export and postprocessing of image files using the Operetta flatfield export script[80] and the Operetta importer plugin[81] in FIJI. Next, several in-house FIJI macros written by the BioImaging and OpticsPlatform (BIOP) at EPFL were used for data analysis. First, image size was further reduced by scaling with a factor of 0.5. Drift correction with the HyperStackReg plugin was performed and the edge of the images was cropped by 40 pixels corresponding to 80 µm. The TrackMate plugin was used for automated cell tracking. First, macrophages were detected based on the CellTracker green channel using the LoG detector (10 µm^2^ particle size, quality threshold of 15). Next, cells were tracked with the SparseLAPtracker (15 µm linking maximum distance, 15 µm gap closing maximum distance, maximum frame gap of 2). Lastly, macrophages were classified as dead or alive based on the DRAQ7 signal. To determine if a cell was dead or alive, the average DRAQ7 background signal was calculated based on the first timeframe. An intensity threshold of 300 was defined, identifying all cells as dead that were 300 intensity units above the background signal. Death was classified by tracks meaning once a track was above the threshold it remained classified as dead. Data was plotted in Python. The percentage of macrophage survival was calculated as:

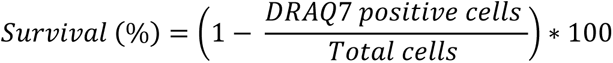

To estimate total survival over 10 h, the area under the curve (AUC) of survival curves for all bacterial strains including noninfected cells (control) was calculated and total survival was estimated as:

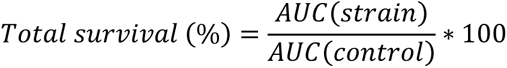

### Statistical testing

All statistical tests were performed in Python except for the Tn-seq data analysis which was conducted in TRANSIT. Typically, a one-way ANOVA with Dunnett’s post-hoc test was applied comparing different mutant strains to a control strain, as indicated in the figure descriptions.

## Acknowledgements

We acknowledge Nicolas Chiaruttini and Olivier Burri from the EPFL Bioimaging and Optics Core Facility (BIOP) for developing the image analysis workflow of the cytotoxicity assay as well as the Lausanne Genomic Technologies Facility for sequencing. Furthermore, we would like to thank Laure Le Blanc for providing guidance in data analysis.

## Author contributions

Conceptualization: T.D. and A.P.

Data curation: T.D.

Formal analysis: T.D. and A.W.

Funding acquisition: A.P.

Investigation: T.D. and C.T.

Methodology: T.D., L.A.M., C.T., Z.A.

Project administration: T.D. and A.P.

Supervision: A.P.

Visualization: T.D.

Writing-original draft: T.D. and A.P.

Writing-review and editing: T.D., L.A.M., R.A.F., A.P.

## Supporting information captions

### Supplementary figures

**Figure S1:** Phagocytosis of the avirulent mutant Δ*popB* and the Tn library in THP-1 macrophages.

**Figure S2:** Gene ontology analysis of intracellular fitness determinants.

**Figure S3:** Phagocytosis imaging assay with *P. aeruginosa* motility mutants.

**Figure S4:** Duration of bacterial contacts with AML cells.

**Figure S5:** Bacterial displacement during macrophage attachment.

**Figure S6:** *P. aeruginosa* cytotoxicity in AML cells.

**Figure S7:** Swimming and twitching motility of clinical isolates.

**Figure S8:** Growth curves of clinical isolates.

### Tables

**Table S1:** Bacterial strains.

**Table S2:** Plasmids.

**Table S3:** Tn-seq analysis for comparison #1: extracellular fraction (supernatant) versus inoculum.

**Table S4:** Tn-seq analysis for comparison #2: intracellular fraction (macrophage lysate) versus inoculum.

**Table S5:** Tn-seq results used for the correlation of comparison #1 and #2.

**Table S6:** List of genes used for functional annotation.

**Table S7:** Functional annotation of Tn-seq data.

### Movies

**Movie S1:** THP-1 macrophage survival during infection with wild-type PAO1.

Macrophages were stained with CellTracker (magenta) and infected at MOI 10. Cell death was monitored by supplementation of the live-dead dye DRAQ7 (cyan) to the medium. This timelapse is one of the technical replicates used for quantification in Figure 1b.

**Movie S2:** THP-1 macrophage survival during infection with the mutant Δ*popB*.

Macrophages were stained with CellTracker (magenta) and infected at MOI 10. Cell death was monitored by supplementation of the live-dead dye DRAQ7 (cyan) to the medium. This timelapse is one of the technical replicates used for quantification in Figure 1b.

**Movie S3:** THP-1 macrophage survival during infection with the Tn library.

Macrophages were stained with CellTracker (magenta) and infected at MOI 10. Cell death was monitored by supplementation of the live-dead dye DRAQ7 (cyan) to the medium. This timelapse is one of the technical replicates used for quantification in Figure 1b.

**Movie S4:** AML cells phagocytosing the avirulent mutant Δ*popB*.

AML cells (CellTracker, magenta) were infected with Δ*popB* (mScarlet, white) at MOI 100. Corresponding movie to images in Figure 2a. The image was cropped for visualization purposes.

**Movie S5:** Exemplary timelapse for phagocytosis quantifications.

AML cells (CellTracker, magenta) were infected at MOI 10 with different motility mutants in the Δ*popB* (mScarlet, white) background. In this movie uptake of the control strain Δ*popB* is shown. This timelapse is one of the technical replicates used for quantification in Figure 2c and 4b.

**Movie S6:** Δ*popB* contact dynamics with AML cells.

Bacterial interactions (mScarlet, white) with AML cells (CellTracker, magenta) were recorded at MOI 200. Bacteria were tracked during contact with macrophages (yellow). This timelapse is one of the replicates used for quantifications in Fig. 3c,d; 4d,e; S4; S5. The image was cropped for visualization purposes.

**Movie S7:** Δ*popB*Δ*fliC* contact dynamics with AML cells.

Bacterial interactions (mScarlet, white) with AML cells (CellTracker, magenta) were recorded at MOI 200. Bacteria were tracked during contact with macrophages (yellow). Corresponding timelapse (cropped image) to quantifications in Fig. 3c,d; S4; S5. The image was cropped for visualization purposes.

**Movie S8:** Δ*popB*Δ*motAB* contact dynamics with AML cells.

Bacterial interactions (mScarlet, white) with AML cells (CellTracker, magenta) were recorded at MOI 200. Bacteria were tracked during contact with macrophages (yellow). This timelapse is one of the replicates used for quantifications in Fig. 3c,d; S4; S5. The image was cropped for visualization purposes.

**Movie S9:** Δ*popB*Δ*motCD* contact dynamics with AML cells.

Bacterial interactions (mScarlet, white) with AML cells (CellTracker, magenta) were recorded at MOI 200. Bacteria were tracked during contact with macrophages (yellow). This timelapse is one of the replicates used for quantifications in Fig. 3c,d; S4; S5. The image was cropped for visualization purposes.

**Movie S10:** Δ*popB*Δ*motABCD* contact dynamics with AML cells.

Bacterial interactions (mScarlet, white) with AML cells (CellTracker, magenta) were recorded at MOI 200. Bacteria were tracked during contact with macrophages (yellow). This timelapse is one of the replicates used for quantifications in Fig. 3c,d; S4; S5. The image was cropped for visualization purposes.

**Movie S11:** Δ*popB*Δ*pilA* contact dynamics with AML cells.

Bacterial interactions (mScarlet, white) with AML cells (CellTracker, magenta) were recorded at MOI 200. Bacteria were tracked during contact with macrophages (yellow). This timelapse is one of the replicates used for quantifications in Fig. 4d,e; S4; S5. The image was cropped for visualization purposes.

**Movie S12:** Δ*popB*Δ*pilT* contact dynamics with AML cells.

Bacterial interactions (mScarlet, white) with AML cells (CellTracker, magenta) were recorded at MOI 200. Bacteria were tracked during contact with macrophages (yellow). This timelapse is one of the replicates used for quantifications in Fig. 4d,e; S4; S5. The image was cropped for visualization purposes.

**Movie S13:** AML cell survival during infection with wild-type PAO1.

Macrophages were stained with CellTracker (magenta) and infected at MOI 10. Cell death was monitored by supplementation of the live-dead dye DRAQ7 (cyan) to the medium. This timelapse is one of the technical replicates used for quantification in Figure 3e,f; 6e,f; S6b,c,d.

**Movie S14:** AML cell survival during infection with mutant Δ*popB*.

Macrophages were stained with CellTracker (magenta) and infected at MOI 10. Cell death was monitored by supplementation of the live-dead dye DRAQ7 (cyan) to the medium. This timelapse is one of the technical replicates used for quantification in Figure S6b.

**Movie S15:** AML cell survival during infection with mutant Δ*fliC*Δ*pilT*.

Macrophages were stained with CellTracker (magenta) and infected at MOI 10. Cell death was monitored by supplementation of the live-dead dye DRAQ7 (cyan) to the medium. This timelapse is one of the technical replicates used for quantification in Figure 5e,f.

