## Supplementary_FigS1-S8_TableS1-S2 for "*Pseudomonas aeruginosa* balances cytotoxicity and motility to counter phagocytosis by macrophages"

**This document contains**

Supplementary Figures S1-S8

Supplementary Tables S1-S2

### Supplementary figures

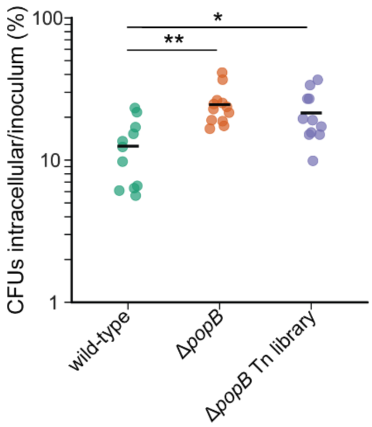


**Figure S1:** Phagocytosis of the avirulent mutant Δ*popB* and the Tn library in THP-1 macrophages.

Intracellular CFUs were quantified after infection of THP-1 macrophages at MOI 10 and gentamycin selection. The uptake of the T3SS mutant Δ*popB* and the Tn library generated in the same background was compared to PAO1 wild-type. Black line, mean; circles, technical replicates (from 3 independent experiments). Statistics were assessed by one-way ANOVA with Dunnett’s post-hoc test.

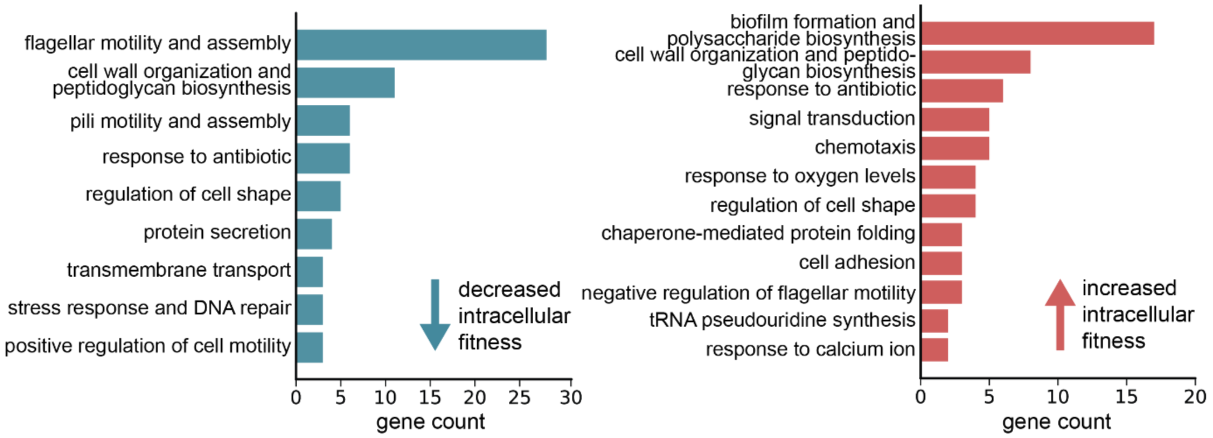


**Figure S2:** Gene ontology analysis of intracellular fitness determinants.

Gene ontology analysis was performed on genes with significant fold changes (adjusted p-value<0.05) in the intracellular fraction of the Tn-seq data set. Results are shown for transposon mutants that were either depleted (blue) or enriched (red) intracellularly.

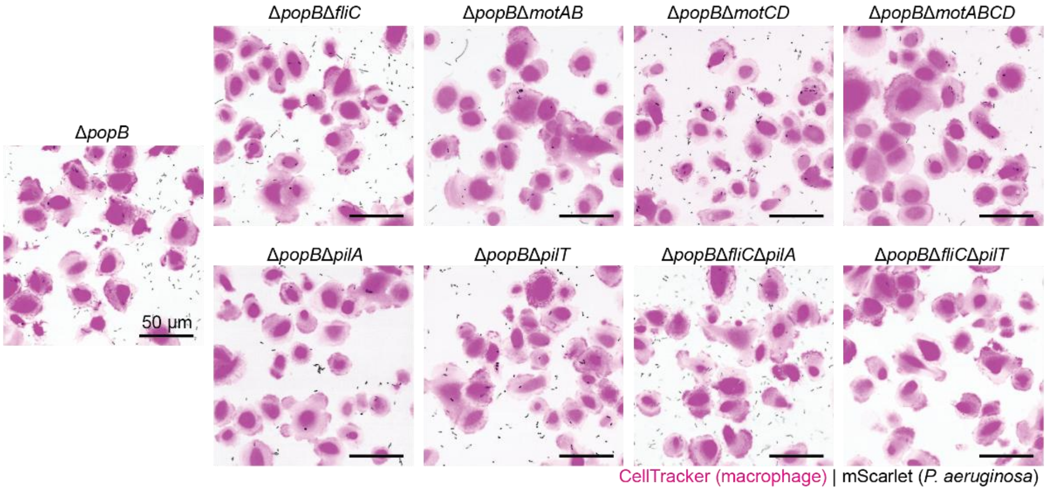


**Figure S3:** Phagocytosis imaging assay with *P. aeruginosa* motility mutants.

Exemplary images of the phagocytosis assay with different PAO1 motility mutants at the imaging endpoint (60 min p.i., first z-plane, cropped images). AML cells were stained with CellTracker and infected at MOI 10 with bacterial strains expressing the fluorescence reporter mScarlet.

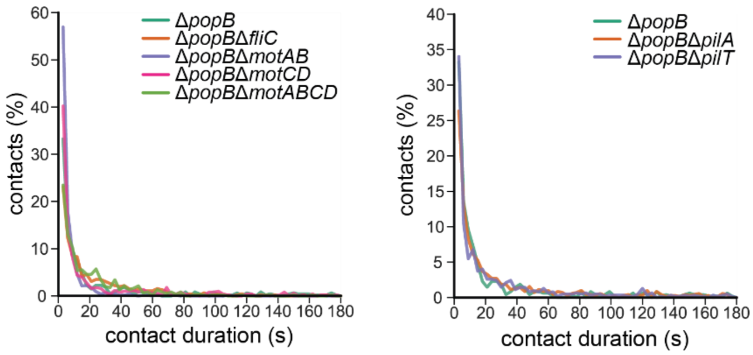


**Figure S4:** Duration of bacterial contacts with AML cells.

AML cells were infected at MOI 200 with flagellar and T4P mutants. Short timelapse movies of 3 min were acquired at a 3 s interval in a single z-plane above the surface. The distribution of contact durations per mutant was plotted. Contact durations were pooled together from all replicates per mutant (n=9).

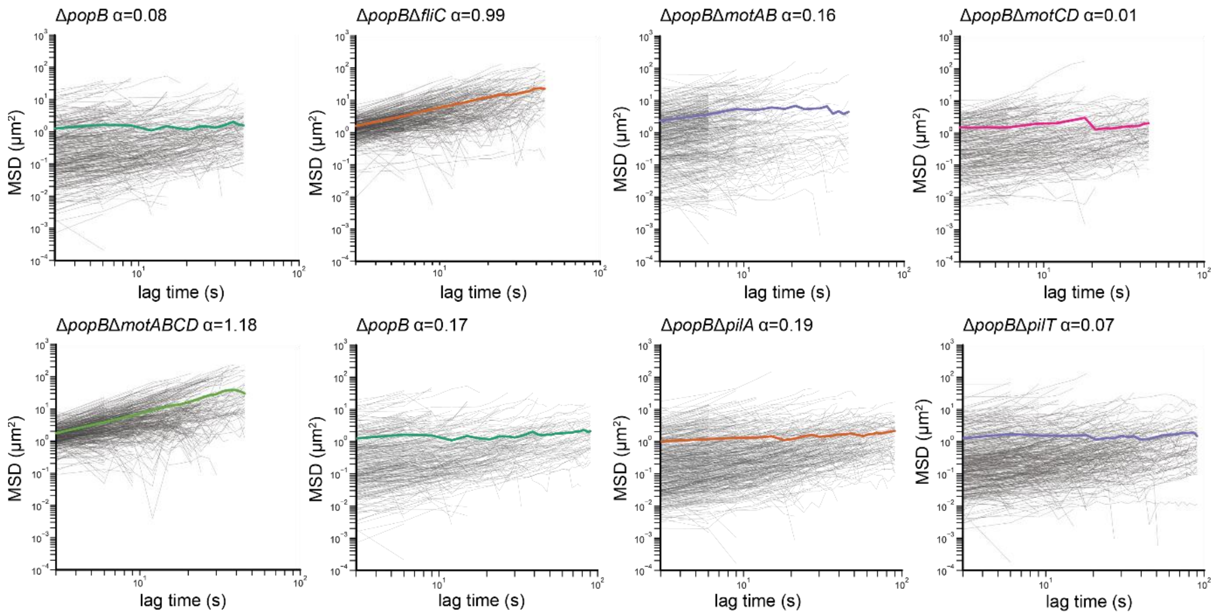


**Figure S5:** Bacterial displacement during macrophage attachment.

The mean square displacement (MSD) of bacteria during contact with macrophages was calculated. Tracking data for all replicates per mutant (n=9) were pooled together. Since contacts vary in time, the tracks have different lengths. The average MSD and individual track MSDs are displayed up to the cut-off, beyond which data sparsity leads to non-linear behaviour in the average MSD in at least one mutant per group (cut-off: 45 s, flagellar mutants; 90 s, T4P mutants). Thick coloured line, mean; thin grey lines, individual tracks; α, diffusion exponent.

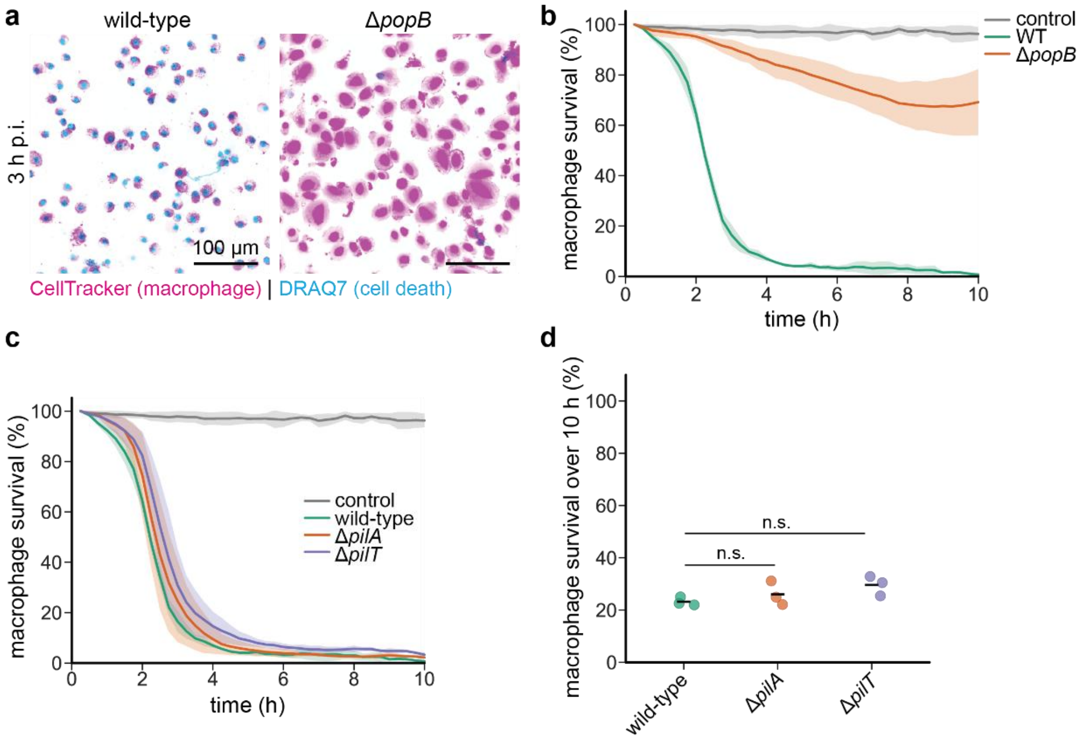


**Figure S6:** *P. aeruginosa* cytotoxicity in AML cells.

AML cells were infected at MOI 10 and macrophage survival was observed over 10 h. Macrophages were stained with the live-cell dye CellTracker. DRAQ7 was added to the culture medium staining the macrophage nuclei upon cell death. **a**, Maximum z-projections (4 planes, 2 µm step size) showing macrophage cell death upon *P. aeruginosa* infection. PAO1 wild-type kills AML cells within 3 h while the mutation Δ*popB* abolishing T3SS is protective. **b**, Quantification of macrophage survival in PAO1 wild-type and Δ*popB*. Uninfected AML cells served as a control. Thick line, mean; shaded area, standard deviation across biological replicates (n=3). **c**, Quantification of AML cell survival when infected with T4P mutants. Thick line, mean; shaded area, standard deviation across biological replicates (n=3). **f**, Total survival over 10 h was calculated as the percentage of the bacterial strain AUC relative to the control AUC (uninfected cells). Black line, mean; circles, biological replicates; statistical test, one-way ANOVA with Dunnett's post-hoc.

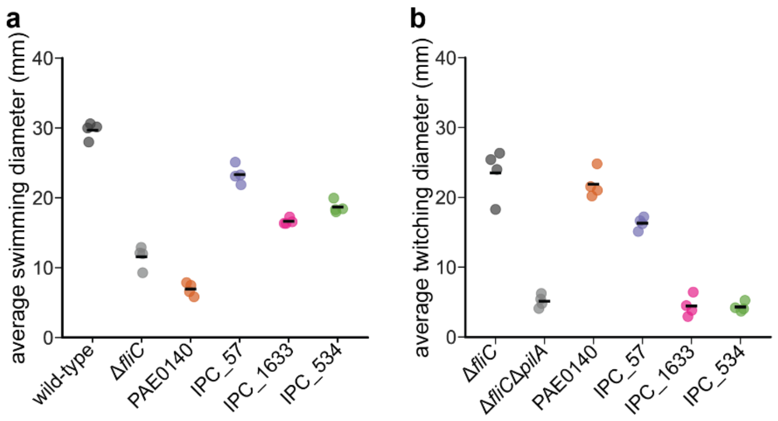


**Figure S7:** Swimming and twitching motility of clinical isolates.

**a**, Swimming motility was assessed by soft-agar plate assay. PAO1 wild-type and Δ*fliC* served as positive and negative controls, respectively. Quantifications of swimming diameters are shown. Black line, mean; circles, technical replicates (two independent experiments). **b**, Twitching motility was measured by stab assay. Non-swimming mutants were used as controls with Δ*fliC* exerting normal twitching motility, and Δ*fliC*Δ*pilA* being unable to twitch. Quantifications of twitching diameters are presented. Black line, mean; circles, technical replicates (two independent experiments).

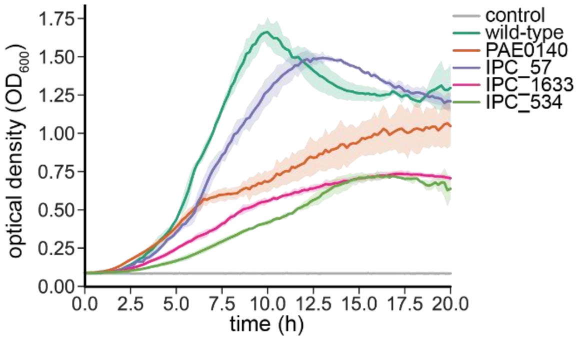


**Figure S8:** Growth curves of clinical isolates.

Clinical isolates and PAO1 wild-type were grown in LB medium over 20 h with plain LB medium serving as a control. The increase in optical density over time is depicted. Thick line, mean; shaded area, standard deviation across biological replicates (n=3).

### Supplementary tables

**Table S1:** Bacterial strains.

| Name and genotype | Source/ Reference | Identifier |
|---|---|---|
| *Pseudomonas aeruginosa* PAO1 | [1] | ATCC 15692 |
| *Escherichia coli* DH5α (hsdR rec lacZYA Φ80 lacZM15) | Invitrogen | n.a. |
| *Escherichia coli* S17.1 (thi pro hsdR recA RP4-2 (Tc::Mu) (Km::Tn7)) | Stratagene | n.a. |
| PAO1 Δ*fliC* | [2] | 177 |
| PAO1 Δ*fliC*Δ*pilT* | this study | 335 |
| PAO1 Δ*fliC*Δ*pilA* | this study | 1047 |
| PAO1 Δ*pilA* | this study | 1208 |
| PAO1 Δ*pilT* | this study | 1269 |
| PAO1 Δ*popB* | this study | 1739 |
| PAO1 Δ*popB*Δ*pilA* | this study | 2207 |
| PAO1 Δ*popB*Δ*pilT* | this study | 2208 |
| PAO1 Δ*popB*Δ*fliC* | this study | 2220 |
| PAO1 Δ*popB*Δ*fliC*Δ*pilA* | this study | 2240 |
| PAO1 Δ*popB*Δ*fliC*Δ*pilT* | this study | 2241 |
| PAO1 Δ*popB*Δ*motAB* | this study | 2604 |
| PAO1 Δ*popB*Δ*motCD* | this study | 2605 |
| PAO1 Δ*motAB* | this study | 2652 |
| PAO1 Δ*motCD* | this study | 2653 |
| PAO1 Δ*popB*Δ*motAB*Δ*motCD* | this study | 2654 |
| PAO1 Δ*motAB*Δ*motCD* | this study | 2656 |

| | | |
|---|---|---|
| Clinical isolate PAE0140 | [3] | 2907 |
| Clinical isolate IPC_57 | [3] | 2932 |
| Clinical isolate IPC_1633 | [3] | 2934 |
| Clinical isolate IPC_534 | [3] | 2935 |

**Table S2:** Plasmids.

| **Name and relevant information** | **Source/ Reference** |
|---|---|
| pEX18AP (Suicide vector based on pUC18, AmpR, ColE1 ori (*E. coli*), oriT, sacB, lacZα) | [4] |
| pEX18GM (Suicide vector based on pUC18, GmR, ColE1 ori (*E. coli*), oriT, sacB, lacZα) | [4] |
| pTNS2 | Addgene 64968 |
| pUC18T-mini-Tn7T-Gm- Ptet_mScarlet | Addgene 63121 (with Ptet promoter fused to mScarlet |
